# How adaptation, training, and customization contribute to benefits from exoskeleton assistance

**DOI:** 10.1101/2021.04.25.440289

**Authors:** Katherine L. Poggensee, Steven H. Collins

## Abstract

Exoskeletons can enhance human mobility, but we still know little about why they are effective. For example, we do not know the relative importance of training, how much is required, or what type is most effective; how people adapt with the device; or the relative benefits of customizing assistance. We conducted experiments in which naïve users learned to walk with ankle exoskeletons under one of three training regimens characterized by different levels of variation in device behavior. Assistance was also customized for one group. Following moderate-variation training, the benefits of customized assistance were large; metabolic rate was reduced by 39% compared to walking with the exoskeleton turned off. Training contributed about half of this benefit and customization about one quarter; a generic controller reduced energy cost by 10% before training and 31% afterwards. Training required much more exposure than typical of exoskeleton studies, about 109 minutes of assisted walking. Type of training also had a strong effect; the low-variation group required twice as long as the moderate-variation group to become expert, while the high-variation group never acquired this level of expertise. Curiously, all users adapted in a way that resulted in less mechanical power from the exoskeleton as they gained expertise. Customizing assistance required less time than training for all parameters except peak torque magnitude, which grew slowly over the study, suggesting a longer time-scale adaptation in the person. These results underscore the importance of training to the benefits of exoskeleton assistance and suggest the topic deserves more attention.

## Introduction

Exoskeletons can make walking easier. For people with movement disorders, exoskeletons can assist gait to overcome impairments. Exoskeletons can help people with cerebral palsy decrease crouch gait (*1*) and energy cost (*2*). People with stroke have demonstrated improvements in energy economy (*3*) and other functional measures (*4*) with the aid of exoskeletons. Exoskeletons can also make difficult tasks easier, such as carrying heavy loads (*5*), walking up inclines (*6*), and running (*7, 8*). Exoskeletons could also overcome some of the effects of aging, for example by decreasing the energy cost of walking in an elderly population (*9*).

Despite the advances in exoskeleton designs, there are still gaps in understanding how humans benefit from exoskeleton assistance. Because there is a human in parallel with the exoskeleton, device performance is necessarily tied to human performance. To understand how humans can master exoskeleton performance, we need to determine not only what optimal human performance looks like but also how to guide people to that optimal gait (*10*).

While aspects of the biomechanical response to exoskeleton use have been characterized in several contexts (*11–14*), little is known about how people adapt to the device. People can modulate their muscle activity to control an exoskeleton with a fixed (*15*) or adaptive controller (*16*), eventually plateauing at a reduced level of activity. While metabolic cost can also stabilize after exposure to a constant exoskeleton controller (*15, 17, 18*), motor learning research suggests that variable training may result in better outcomes (*19–21*). Estimates of the amount of time required to reach a steady-state minimum energy cost ranges widely in the literature, from 18 (*17*) to 90 minutes (*15*), but there are indications among these studies that longer exposure may have resulted in greater benefits.

In addition to the improvements possible with variation training, there is evidence to suggest that subject-specific controllers may result in better exoskeleton assistance. Human gait is so personalized that it can be used as a biometric measurement (*22*). Customization in exoskeleton control can exploit these variations (*23*) and has been shown to result in large reductions in metabolic cost with exoskeletons (*24, 25*). Despite its success, there have been no studies to determine the relative importance of a personalized controller.

We designed an experiment to understand how the type and duration of training affects performance for both fixed and personalized exoskeleton controllers. Participants were trained in bilateral ankle exoskeleton walking for nearly ten hours, distributed across one pre-test day and five days of training. We demonstrated that human-in-the-loop optimization can simultaneously train exoskeleton users and discover customized assistance profiles. Motor adaptation accounted for half of the reduction in energy cost, highlighting the importance of fully training participants before evaluating performance of an assistive device.

## Results and Discussion

Participants with no prior assistive device experience were exposed to exoskeleton assistance to determine how different types of training and customization affect performance (Fig 1). A pre-test was conducted on the first day to determine how people initially responded to the exoskeleton in a fixed, generic assistance pattern. The generic profile was the average optimized profile from a two-day pilot experiment with ten participants (Supplementary Materials). Performance with assistance was compared to baseline trials with an unpowered, zero-torque mode and with normal running shoes. Each subsequent day began with a 72-minute adaptation trial, determined by training group. Adaptation was followed by a series of validation tests, which included the conditions tested in the pre-test as well as customized profiles for some participants.

**Figure 1:**
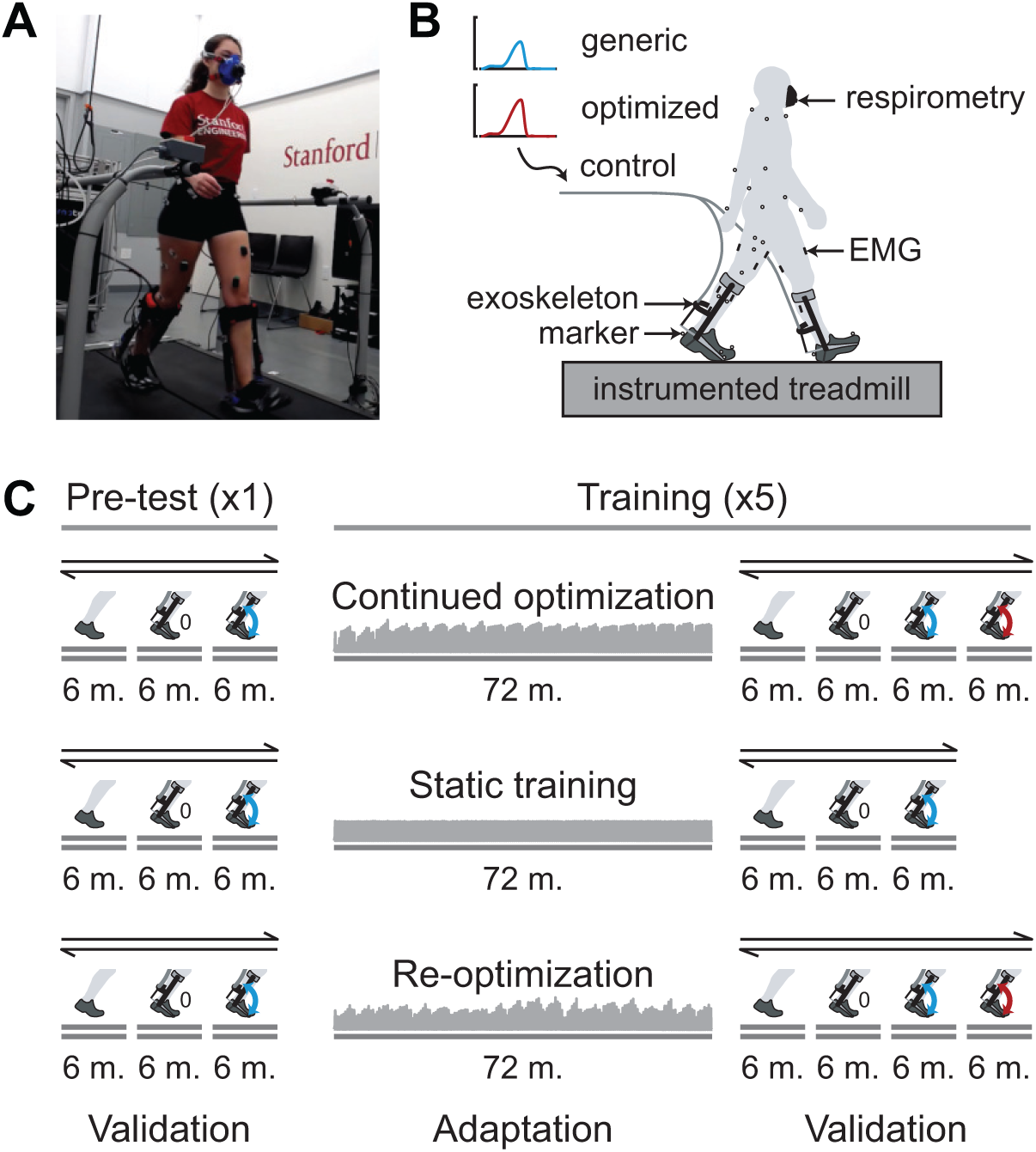
Experimental setup and protocol. (*A*) Photograph of the experimental setup. Participants wore tethered bilateral ankle exoskeletons and walked on an instrumented treadmill. (*B*) Schematic of the experimental setup. Metabolic energy consumption, muscle activity, ground reaction forces, and body kinematics were recorded throughout the experiment. The devices were controlled by off-board motors. (*C*) Schematic of the experimental protocol. Each row represents the protocol for a typical participant in each of the three training groups. Participants began with a pre-test of six six-minute validation trials: two trials wearing running shoes and no exoskeleton, two trials with the exoskeleton unpowered in a zero-torque condition, and two trials with a generic assistance profile. On each of the five training days, participants began with an adaptation trial, which differed by training group. All training groups then experienced validation trials, including the six trials from the pre-test day; the *continued optimization* and *re-optimization* training groups also experienced customized assistance for two validation trials. The validation tests were randomized and presented in a double-reversal ABCDDCBA order to mitigate the effects of noise in the metabolics measurements and trial order.

The fifteen participants were randomly sorted into three training groups, two of which experienced human-in-the-loop optimization. During optimization, a series of assistance profiles were randomly sampled based on the estimated optimal profile. Participants experienced each of these profiles for two minutes, and their energy cost was estimated for each. The algorithm then ranked the assistance profiles by energy cost to determine a new estimate for the optimal assistance profile.

The *continued optimization* group underwent human-in-the-loop optimization throughout the adaptation period. The optimization was seeded with a comfortable assistance profile and a large distribution from which to sample control laws; these settings have been shown to result in beneficial assistance (*24*). On subsequent days, the optimization began with the estimated optimal profile and associated distribution from the previous day. The algorithm sampled from a narrower distribution as the confidence in the optimal estimate increased, so this training program began with variation training before converging on profiles near the optimum. On each day of testing, the estimated optimal profile from the end of that day was included in the validation tests; the customized profile on day *n* was then the result of 72*n* minutes of optimization. The generic profile was periodically tested throughout the adaptation trial to track adaptation.

The *static training* group only experienced generic assistance for the duration of the adaptation period. This training regimen was intended to isolate the effects of training time on user expertise. Training protocols with fixed exoskeleton behavior are the most common in prior studies (*15, 17, 18*).

The *re-optimization* training group underwent human-in-the-loop optimization as described above, but with the same initial seed used on each day. Because there was less time for the optimizer to converge, this training program had higher variability than either the *continued optimization* or the *static training* group. The customized profiles were determined only by the 72-minute trial on that day rather than a cumulative estimate. Generic assistance was also periodically applied to track motor adaptation.

### Training resulted in a substantial improvement in metabolic energy consumption

With only 12 minutes of lifetime exposure to exoskeleton assistance by the end of the pre-test day, most participants exhibited reduced metabolic cost with generic assistance, with an average improvement of 10.0 ± 11.0% (mean ± standard deviation) relative to zero torque (Fig 2A, paired *t*-test, *P* = 0.009). After at least four hours of training, participants received a larger benefit from generic assistance (Fig 2B). Once fully adapted, the *continued optimization* group experienced an energy cost reduction of 30.6±8.1% relative to the zero torque condition (paired *t*-test, *P* = 0.005), and the *static training* group reduced their energy cost by 28.2±4.0% (paired *t*-test, *P* = 2.74*e* − 5). The *re-optimization* group did not improve their energy economy to the same degree as the other two groups, but still experienced a larger improvement than the average pre-test response: by 17.5 ± 9.2% compared to the zero-torque condition (paired *t*-test, *P* = 0.09).

**Figure 2:**
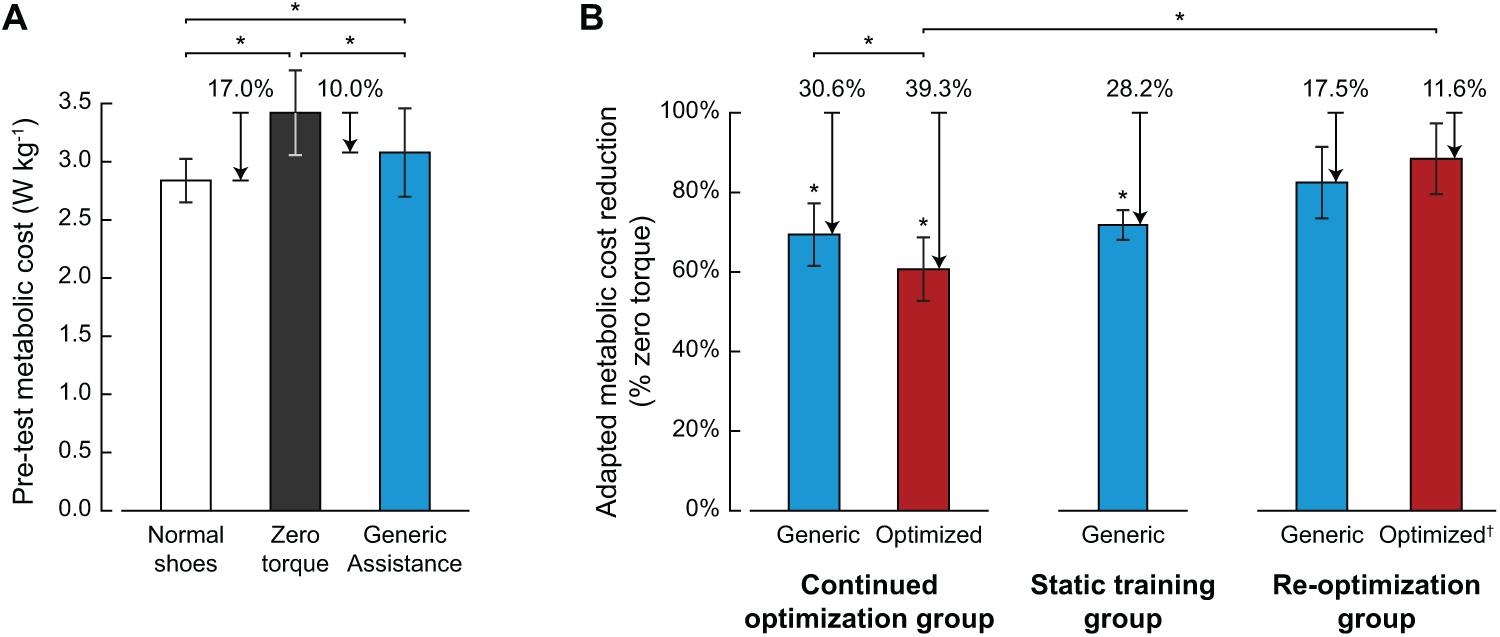
Metabolic cost with assistance during (*A*) the pre-test and (*B*) after adaptation. (*A*) Metabolic cost (W·kg^−1^) during the normal shoes, zero torque, and generic assistance conditions for all participants. Initially participants showed a small, but statistically significant, reduction in metabolic cost in the generic assistance condition compared to the zero torque condition. (*B*) Metabolic cost of the generic and optimized assistance conditions, normalized to the zero torque condition on that day, for the three training groups. All conditions at the end of the experiment were significantly different than zero torque. Optimized assistance for the *re-optimization* group (†) is the response to the torque profile produced by the optimization algorithm at the end of that day; it is not assumed to be optimally assistive. Comparisons within training groups were made using paired *t*-tests. Comparisons across training groups were made using unpaired *t*-tests. Error bars denote standard deviation. Statistical significance between conditions is denoted with an asterisk for *P <* 0.05; asterisks above a single bar denote significance with respect to zero torque.

We calculated the change in metabolic cost relative to the energy cost of walking with zero torque in order to isolate the benefits of assistance from the device-specific costs of added leg mass or volume and the day-to-day fluctuations of metabolic cost seen in normal walking (*26*). The experiment was performed on a versatile emulator system (*27*), designed to be powerful and modular with off-board motors and generic end-effectors. An untethered version could be more streamlined, given a specialized function, or bulkier, because actuation would need to be on-board. Adaptation to zero torque was fast (27 minutes, Supplementary Materials), but with some intersubject variability. The final, adapted results were not affected, but may have resulted in a higher baseline value during the pre-test, meaning that the training effect we identified is a lower bound.

Participants reduced their metabolic cost almost immediately upon donning a powered exoskeleton for the first time, but tripled the benefit from assistance with more training. While some participants increased their energy cost on the first day of testing, the average response is comparable to other single-session studies (*5, 28*). Many participants initially disliked assistance, but improved their opinions by the end of the experiment. We did not directly measure subjective preferences, which may give more insight into any correlations between metabolic cost and preference (*29*). Prior to testing, we did not familiarize participants with assistance in order to fully track their motor adaptation. Many exoskeleton studies report habituating participants before measuring their responses (*28*). Training may have varying effects on energy economy, depending on the mode of assistance or the complexity of the task, but these results support the general need for sufficient training to see larger reductions in metabolic cost.

### Metabolic cost exponentially decreased with training

The generic assistance profile was experienced at least thirty-two times throughout the experiment in two- and six-minute trials. The performance on each trial was used to track adaptation as a function of the time a participant was exposed to exoskeleton assistance. At the study level, metabolic cost decreased over time, following an exponential decay of the form

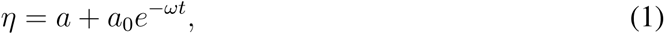

where *η* is the metabolic energy cost, *t* is exposure time in minutes, *a* is the steady-state metabolic cost, *a* + *a*_0_ is the initial energy cost, and *ω* is a time constant (Fig 3A). This exponential model fit the data better than a constant model (ANOVA, *P <* 2.2*e* − 16).

**Figure 3:**
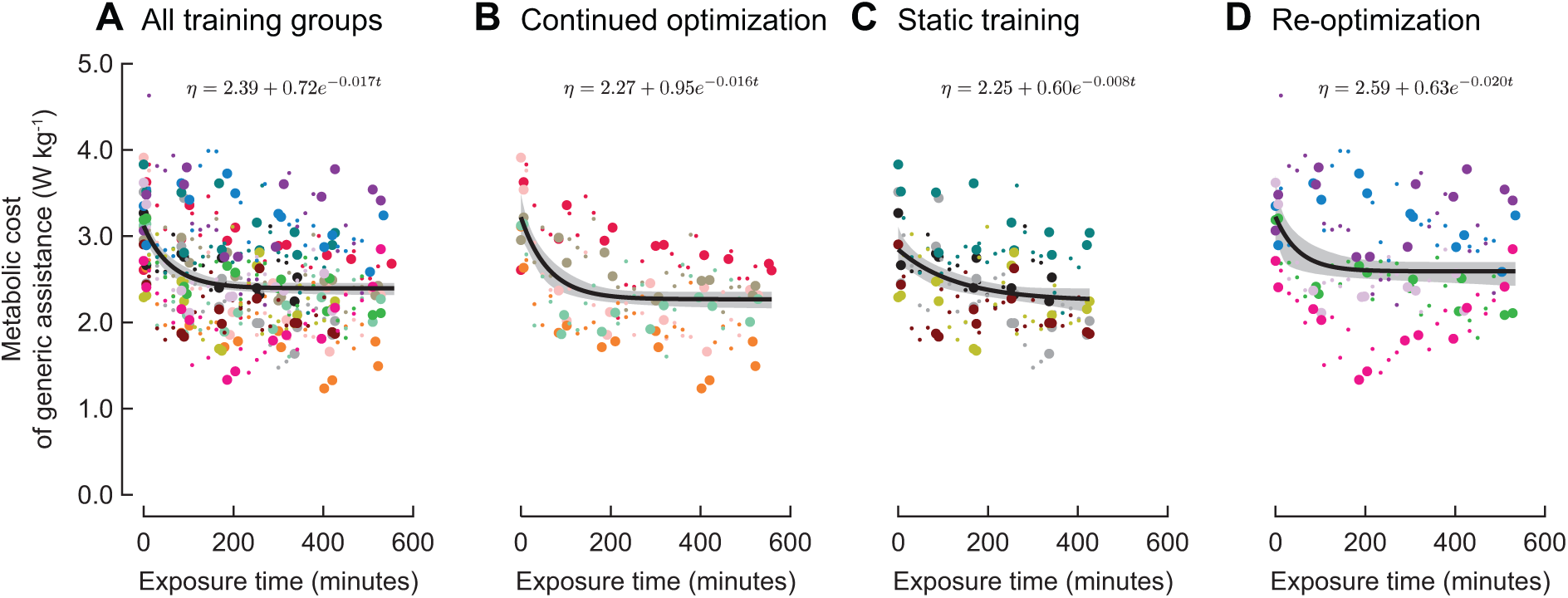
Metabolic cost of generic assistance. The metabolic cost of the fixed generic assistance conditions was modeled as an exponential function of time at (*A*) the study level and at training group levels for the (*B*) *continued optimization*, (*C*) *static training*, and (*D*) *re-optimization* training groups. At the study level, participants need nearly two hours of exposure to active exoskeleton assistance in order to reach expertise, but the rate and expected metabolic reduction depends on the type of training experienced. Individual participants are represented by distinct colors, and 6-minute validation trials are represented by larger points than the 2-minute estimates taken during the adaptation trials. The exponential models, written at the top of each plot, are shown in black with corresponding 95% confidence bands in gray.

We considered several factors before converging on an exponential model of metabolic cost. There are many accepted methods and forms for modeling motor adaptation (*19, 30–34*). The aspects of exoskeleton walking that guided our choice of model are outlined here; an extensive comparison of models can be found in the Supplementary Materials. The chosen exponential function, shown in Eq. 1, is often used to describe motor learning in locomotion research (*15, 17, 34, 35*). In order to isolate the effects of motor adaptation, we modeled only the responses to a fixed exoskeleton assistance pattern, in this case the generic profile.

We used metabolic cost as a proxy for the learned control pattern. Metabolic cost is an important outcome in exoskeleton research (*10*) and other locomotion-based motor learning research (*36, 37*). Kinematic measures are often used in learning research (*38, 39*), but are thought to stabilize earlier than metabolic cost (*40*). We would then expect models using metabolic cost to be the most conservative for determining the time to expertise.

Metabolic measurements were modeled with respect to assisted walking exposure time and were averaged within a single two- or six-minute trial. Developmental research suggests that time spent walking is a better indicator of children’s ability to walk than days since onset of walking (*41*). Exoskeleton learning literature has reported adaptation as a function of single-breath data (*17*) and averaged metabolic cost (*18*). The dynamics of metabolic cost at the beginning of a walking trial (*42*) confound exponential models by artificially exaggerating the warm-up period described in other motor learning paradigms (*43, 44*). While data averaging can skew results if there is training that occurs during the period of time over which the data are averaged (*39*), the trials were short enough in this case to not encode substantial adaptation, and the averaged values carry greater meaning. This analysis was insensitive to factors such as the number of rest days and data averaging over individual trials, which are explored in the Supplementary Materials. Overall, the modeling methods presented here characterized the learning process well and can be used to track adaptation to similar assistive devices.

### Full adaptation can be a slow process

We defined time to reach expertise as the time before the model fit of metabolic rate was within 5% of its asymptote. At the study level, subjects needed 109 minutes of practice (IQR, 91 to 130 minutes) to become expert exoskeleton users. This is significantly longer than the length of most exoskeleton studies, including those for which adaptation time is the main result (*15, 17, 18*).

This experiment was long enough to see steady-state behavior over an extended period of time, increasing our confidence in the final measurements compared to a shorter protocol (*36, 45*). If there is an insufficient amount of data at steady state, an exponential model will tend to underestimate the asymptote and consequently underestimate the time for the predicted model to reach that asymptote (*43*). Premature analysis of motor learning in exoskeleton walking may then explain the shorter adaptation times seen in the literature (*17, 18*). Many existing exoskeleton studies incorporate some level of habituation before the start of the experiment without indication that participants are fully trained (*8, 28*). Ensuring participants are fully trained should be an important factor in designing assistive device experiments in the future.

Adaptation time, like any other motor learning outcome, is highly context-dependent (*46*), so we would expect that the exposure time necessary to gain expertise in using an assistive device may vary widely. We may predict that unilateral devices (*15*), devices that assist dorsiflexion, or devices that act at proximal joints (*47, 48*) are easier to master than the bilateral ankle plantarflexor exoskeletons used here. Conversely, increasing the number of joints may increase the complexity of the motor learning problem (*49*) as people often freeze joints (*50*) or decompose movements by joint (*51*) as they learn complex, multi-joint movements, resulting in longer adaptation times. We would expect adaptation time to change as a function of assistance magnitude. For example, the generic assistance used in this study is similar in magnitude to other devices used for augmentation (*5,12,15,16,52*), but higher levels of assistance may be necessary to see benefits in strenuous tasks such as running (*8*). Although devices used to assist populations impaired by neurological disease might benefit from lower levels of assistive torque, motor learning is often impaired in these populations (*53, 54*), so these populations would either need more time to learn or more targeted training. Training methods such as biofeedback (*51, 55*) could speed motor learning, but care should be taken to implement feedback that does not inhibit learning in the target population (*56*). Extensively documenting adaptation times for every type of assistive device may be infeasible, so researchers should consider the implications of training when designing their protocols, either by building training into the experiment or by characterizing adaptation to better predict the steady-state response to assistance.

The burden of training participants may change how exoskeleton research is done in the future. Recruiting participants is a difficult task, and longer duration studies are at increased risk of participant dropout (*57*). Having a pool of trained exoskeleton users rather than sampling from novices may facilitate research on exoskeleton design and control. Studies with smaller sample sizes would allow for more time and resources to be spent on training, enabling researchers to answer more difficult questions on how assistive devices impact human movement (*58*). While these issues are of less concern for people using assistive devices in daily life, where high exposure times are natural and assistance profiles may be easier to use than those developed for the lab, these results highlight the importance of longitudinal testing in the development of future commercial devices.

### Appropriate variation training speeds adaptation

The three training groups experienced different levels of variation in assistance profiles and learned how to use the generic assistance over different timescales (Fig 4A). The *static training* group had zero variation and reached expertise in 218 minutes (IQR, 143 to 358 minutes). While the *static training* group had less exposure time to assisted walking compared to the optimization groups (432 minutes compared to 540 minutes), the time to reach expertise was within the exposure time experienced during this experiment. The *continued optimization* group had large variation at the beginning of training, with the total variance in control parameter values being 71 during the first two days, before tapering to minimal variation by the end of training, with a total variance of 33 over the last two days. This appeared to speed training with participants adapting in 131 minutes (IQR, 107 to 160 minutes). The *re-optimization* training group was exposed to a large variety of assistance profiles throughout the experiment with the total variance of control parameters being 93 across the entire training period. The model predicts that their metabolic cost stabilized after 81 minutes (IQR, 51 to 124 minutes). However, the fact that this group did not achieve the same level of energy cost reduction as the *continued optimization* or *static training* groups suggests that the *re-optimization* group did not attain the same level of expertise as the other training groups.

**Figure 4:**
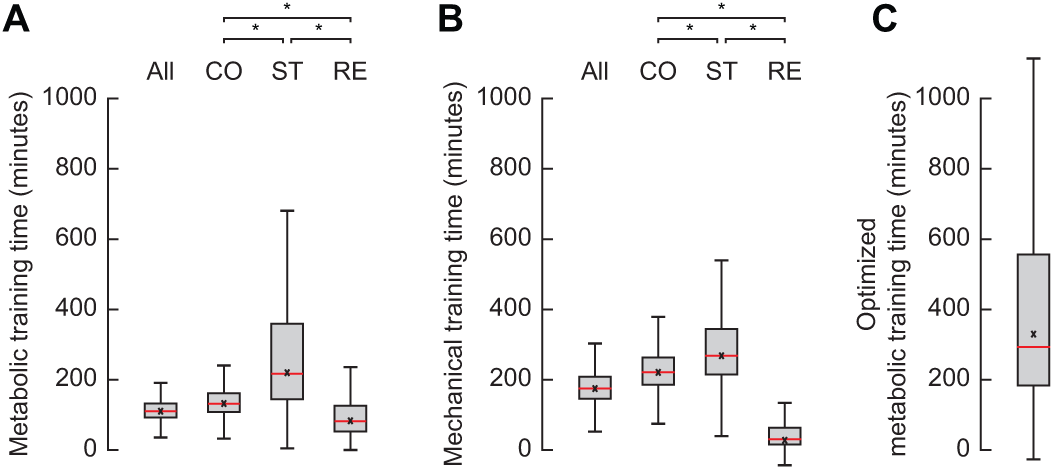
Training time for each training group. (*A*) Training time (defined as the time to reach within 5% of the asymptote) determined by the model of metabolic cost adaptation with generic assistance. (*B*) Training time determined by the model of mechanical power of generic assistance. (*C*) Training time determined by the model of metabolic cost adaptation with optimized assistance. Training time distributions were computed from the bootstrapped models used to compute the confidence bands. The central red line denotes the median, with the top and bottom edges of the box denoting the 75th and 25th percentiles, respectively; the whiskers extend to the most extreme data points not considered outliers. The predicted training time for the models given in the main text are denoted by a black x. Significance between groups is designated with an asterisk (Wilcoxon rank sum test, *P <* 0.05).

The *continued optimization* training protocol appeared to be the most effective. There was a significant improvement in energy consumption after adaptation, compared to the pre-test (paired *t*-test, *P* = 0.005). These participants also adapted more quickly than the participants in the *static training* group (Fig 4A, Wilcoxon rank sum test, *P <* 2.2*e* − 16) and settled on a larger reduction in energy cost than the *re-optimization* training group (Fig 2B, unpaired *t*-test, *P* = 0.04). These results suggest that large variation followed by more targeted practice near the optimized solutions, as provided by the *continued optimization* protocol, may be an effective training approach for exoskeleton studies.

Despite the slower adaptation, simply exposing people to a generically good assistance profile for a long period of time may be as effective in the long term as the training experienced by the *continued optimization* group and more effective than the highly variable training protocol. The *static training* and *continued optimization* groups stabilized to similar reductions in metabolic cost (Fig 2B, unpaired *t*-test, *P* = 0.62), and the predicted steady state energy cost of walking with generic assistance was similar between these two groups, at 2.27 W·kg^−1^ for the *continued optimization* group and 2.25 W·kg^−1^ for the *static training* group. Coincidentally, there were participants in the *static training* group who exhibited large reductions in energy cost in the pre-test before any group-specific training, which may explain why the *static training* protocol did not result in a significant improvement compared to the pre-test (paired *t*-test, *P* = 0.25). Adapted energy savings were consistent with the reductions seen for the *continued optimization* group, so *static training* should still be considered an effective training protocol.

Conversely, the *re-optimization* training protocol was ineffective compared to the other training protocols. The mean energy cost reduction was increased, but results varied widely across participants, and the improvement from the pre-test was not statistically significant (paired *t*-test, *P* = 0.73). By the end of the experiment, some participants, who had showed improvements in the early stages of the experiment, returned to the same performance levels as the pre-test (Fig 3D). These results suggest that participants either were untrained by the end of the experiment or quickly learned a different motor task that appeared to be ineffective by our metrics, demonstrating the trade-off between expertise in a single assistance profile and a coordination pattern that is robust to a wide variety of assistance patterns.

External variability is generally thought to improve motor learning (*19–21, 46, 59, 60*). The variability in movements induced by the exoskeleton assistance may have prompted increased exploration in participants (*46, 60*), resulting in faster adaptation in the *continued optimization* group compared to the *static training* group. The nature of the human-in-the-loop optimization algorithm directed the assistance profiles to converge on those which decrease effort. Moderate rewards can increase participants’ willingness to explore different coordination patterns, thereby speeding learning of an optimal movement pattern (*46*). The similar final performance between the *continued optimization* and *static training* groups are consistent with these ideas. Variability of practice is thought to improve generalizability and long-term retention, but we did not compare these outcomes between groups.

Variability can also interfere with learning, as seen in the *re-optimization* training group. For stable motions, variability can promote exploration along nonoptimal coordination strategies (*60*). The assistance profiles experienced by the *re-optimization* group during training likely provided less reward than the assistance profiles experienced by the *continued optimization* group. Participants may have taken a risk-averse approach and chosen not to explore new movement patterns (*46*), resulting in a single motor control that is effective across all exoskeleton behaviors they experience. While this may be generally acceptable, if the pool of exoskeleton assistance profiles contains detrimental patterns, the universal motor controller may have performed poorly for beneficial exoskeleton behaviors. The level of assistance likely also amplified the unfavorable torque profiles, exacerbating the need for a risk-averse gait. More research should be done to determine when assistance profiles cease to assist and how to appropriately bound variation training.

### Ankle kinematics adapted to exponentially decrease mechanical work derived from the exoskeleton

The generic assistance profile was defined as a fixed time-based torque pattern. Participants were able to modulate their kinematics to determine the amount of mechanical power they received from the device. Interestingly, mechanical power of the fixed assistance pattern decreased in an exponential pattern with training (Fig 5A).

**Figure 5:**
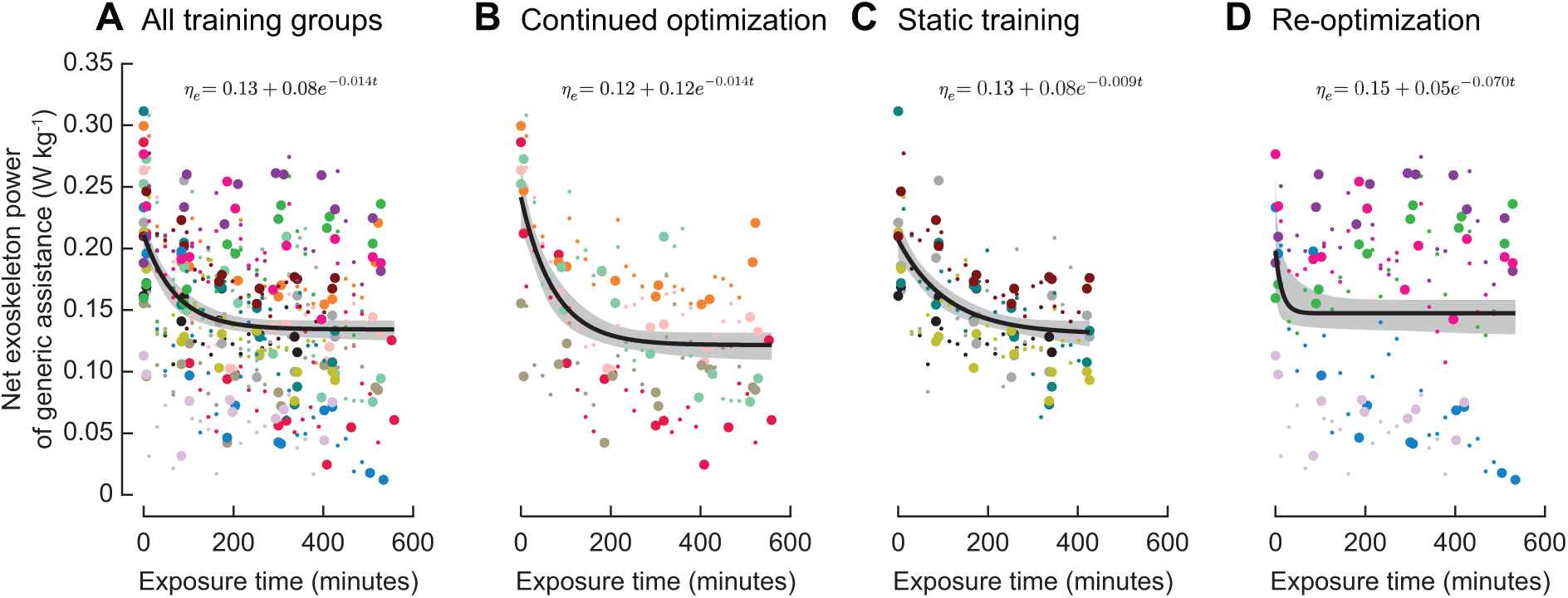
Mechanical power of generic assistance. The mechanical power of the fixed generic assistance conditions was modeled as an exponential function of time at (*A*) the study level and at training group levels for the (*B*) *continued optimization*, (*C*) *static training*, and (*D*) *re-optimization* training groups. Mechanical power decreased with training time at the study level and the group levels. Because the torque profile was fixed in time, the reduction in mechanical power represents kinematic adaptation to the generic assistance. Individual participants are represented by distinct colors, and 6-minute validation trials are represented by larger points than the 2-minute averages taken during the adaptation trials. The exponential models, written at the top of each plot, are shown in black with corresponding 95% confidence bands in gray.

Decreasing plantarflexor muscle activity is a well-documented response to ankle assistance (*61*). There is some evidence that people decrease ankle plantarflexion and exoskeleton mechanical power as well, but only for exoskeletons controlled by muscle activity (*15*). While we did not analyze muscle activity in this paper, we expect the decrease in ankle plantarflexion to be the result of decreasing plantarflexor activity. The exponential decay suggests that there is a neural component to this kinematic adaptation rather than a purely mechanical explanation which would have been immediately evident (*62*).

A prevalent theory in exoskeleton research is that designers should create devices that inject large amounts of mechanical power in order to see larger reductions in metabolic cost (*5*). Previous exoskeleton research has shown that profiles that supply more mechanical work result in larger reductions in energy cost with some evidence of a fixed conversion between external mechanical power and metabolic energy savings (*12, 13*). The adaptation leading to decreased mechanical power and metabolic cost seen in our participants is inconsistent with these models and indicates that there are more complex human dynamics that should be considered when designing exoskeletons.

In other motor learning paradigms such as reaching tasks, kinematics stabilize before metabolic cost (*40*). Mechanical power in our study stabilized in 212 minutes, decreasing at a slower rate than metabolic cost (Fig 4, Wilcoxon rank sum test, *P <* 2.2*e* − 16). Adaptation may follow different dynamics under different contexts. For example, reducing muscle activity may be a higher priority than stabilizing kinematics for long experiments, and the exact durations may vary depending on the type and level of assistance. Metabolic cost alone does not explain choices in gait (*63*), and the factors governing how people explore and adapt are likely similarly nuanced. Identifying other dimensions along which people explore and adapt in response to assistive devices may lead to improved training protocols based on biofeedback and targeted instructions.

### Optimizing assistance resulted in larger energetic benefits

The *continued optimization* group experienced even larger improvements in energy economy when walking with optimized assistance parameters: 39.6±8.2% compared to the 30.6% reduction with generic assistance (Fig 2B, paired *t*-test, *P* = 0.03). The improved energy economy with optimized assistance corroborates the hypothesis that customization, along with a good generic baseline condition and proper training, plays a role in reducing metabolic cost (*24*).

The exact ratio of these factors will necessarily depend on the population and the device. The participants in this study were young, healthy adults with no previous exoskeleton experience nor a history of movement disorders. As exoskeletons become more available, there may be motor learning transfer between different styles of exoskeleton, so training may contribute less to the overall benefit. Diseases such as stroke or surgeries like amputation uniquely affect a person’s mobility and thus customization of assistive devices could be a more important factor. Among the young, healthy population tested in this study, there was a range of energy cost reductions from 25% to 59%. While this large range was unexplained by characteristics such as basic demographic information (Supplementary Materials), there may be other similarities among participants who respond well to assistance (*64*).

While the difference between generic and customized assistance was statistically significant for our study, the real-world significance will depend on context. For homogeneous populations, the extra 10% energy savings may not outweigh the added time and effort needed to optimize assistance. The added complexity in customizing parameters may lead commercial producers to use the generic solution. In heterogeneous populations, some amount of customization may be needed to obtain any benefits.

This human-in-the-loop optimization algorithm yielded large energy benefits, but the difference in energy cost may be attributable to other factors. The optimization was seeded with a different controller than the generic profile, and the resulting mean of the optimized parameters differed from the generic control parameters. We also did not systematically sample the space to determine the sensitivity to changes in parameter values. Future work to determine the importance of slight variations in parameters around a generically beneficial controller may yield finer insights into the implications of customization of device parameters.

### Optimization resulted in parameters that differed from the generic pattern, with some shared characteristics across subjects

We used a genetic algorithm (*24*) to optimize four parameters which determined a smooth, single-peak torque trajectory as a function of stride time. One parameter determined the peak torque magnitude, *τ*, and three parameters determined the timing of the peak, *t_p_*, the rise time, *t_r_*, and the fall time, *t_f_*. After 20 generations, the optimized torque pattern, based on the average of the optimized parameters, was characterized by a peak of 0.68 ± 0.06 N·m·kg^−1^ at 54.3 ± 0.7% stride, which was higher in magnitude and later in time than generic assistance (Fig 6A, paired *t*-test, *P* = 0.007 and *P* = 0.01, respectively). Other aspects of the optimized torque curves, such as the rise time of 27.8 ± 4.5% and fall time of 9.7 ± 0.9%, were similar to generic assistance (Fig 6A, paired *t*-test, *P* = 0.46 and *P* = 0.85, respectively).

**Figure 6:**
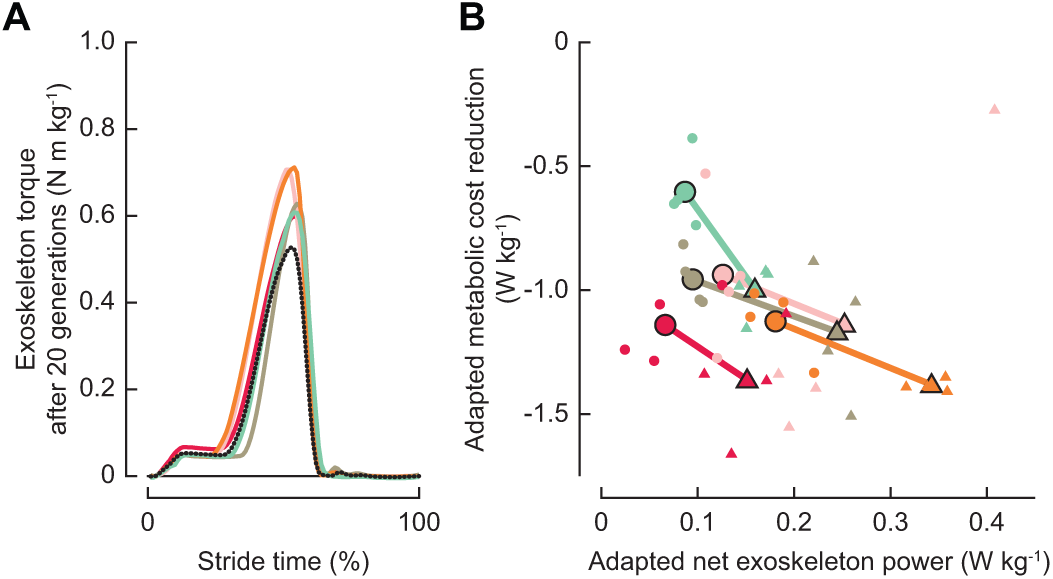
Optimal and generic assistance profiles for the *continued optimization* group. (*A*) Optimal torque trajectories (N m kg^−1^) after twenty generations of optimization, shown in solid lines for each participant. The generic assistance torque trajectory is shown as a dotted line in black. (*B*) Metabolic cost reduction vs. mechanical work for the generic assistance profiles, denoted, and for the optimal assistance profile, denoted. Averaged values are depicted with a black outline. Optimal assistance provided more mechanical work than generic assistance (repeated measures ANOVA, *P* = 2.9*e* − 10).

The average optimized values for peak time and fall time indicate a universally beneficial peak time and offset time for exoskeleton assistance. The peak time and offset time, defined as the sum of the peak time and fall time, were near the limits of the parameter space and did not vary considerably between subjects, with ranges of less than 2% of stride for each node. These limits were chosen based on comfort (*24*), but could be relaxed for different devices. These results, along with results seen during pilot testing and other studies (*8, 24*), indicate that an assistance profile with peak torque near the end of stance, quickly tapering to zero torque at toe-off, results in the largest energy benefits.

Rise time varied substantially across participants, which could be the result of individual variation or the result of a weak relation between rise time and metabolic cost such that rise time does not substantially affect metabolic cost over a large range. It is difficult to determine if rise time is necessarily a subject-specific parameter or if it does not directly affect metabolic cost. To truly understand the importance of rise time as a parameter, another experiment to test energy expenditure in response to a range of rise times, with sufficient training, should be performed.

As participants learn to use the exoskeleton, they benefit more from larger magnitudes of torque assistance. The participants in the pilot test that determined the generic controller would not have fully adapted to exoskeleton assistance because the protocol only lasted two days (Supplementary Materials). This seems to explain the lower magnitude relative to these optimized profiles. Participants in this study initially noted that the level of assistance in the generic profile seemed too high, but were more comfortable by the end of the experiment. This increased comfort could be due to neuromuscular changes, like increased strength at proximal joints, as well as changes in how people perceive assistance timing and magnitude, both of which may be important factors for walking with higher levels of assistance. We expect the optimal peak torque for fully trained users to be much larger in magnitude, based on the trajectory of the parameter during optimization, but within the range that is observed for other ankle exoskeleton devices (*12, 24, 52*). Increased training time or more targeted training may allow people to utilize the larger torques and transfer the mechanical assistance to other joints (*47*). This might happen naturally with commercial or prescribed devices that are used regularly for long periods of time.

### Human-in-the-loop optimization converged over multiple timescales

By the nature of our study, participants co-adapted with the exoskeleton. Therefore, we need to consider both the device parameters and human response in determining the time course of adaptation for the full system. The same exponential model analysis as in Eq. 1 was used to understand the evolution of both the parameters and the metabolic cost of the optimized conditions.

There are three timescales for optimization: convergence time of the timing parameters, convergence time of the peak torque magnitude, and adaptation time of the user. Timing parameters quickly converged, with peak time converging to a value within 5% of the predicted asymptote by 2.0 generations (Fig 7A, IQR, 1.9 to 2.2 generations). Rise time and fall time both converged within a generation, at 0.7 generations for rise time (Fig 7B, IQR, 0.7 to 2.8 generations) and 0.7 generations for fall time (Fig 7C, IQR, 0.6 to 3.4 generations). Peak torque magnitude does not appear to have converged within this experiment, with the model predicting convergence after 106 generations (Fig 7A, IQR, 28 to 108 generations), which is equivalent to 1908 minutes of optimization.

**Figure 7:**
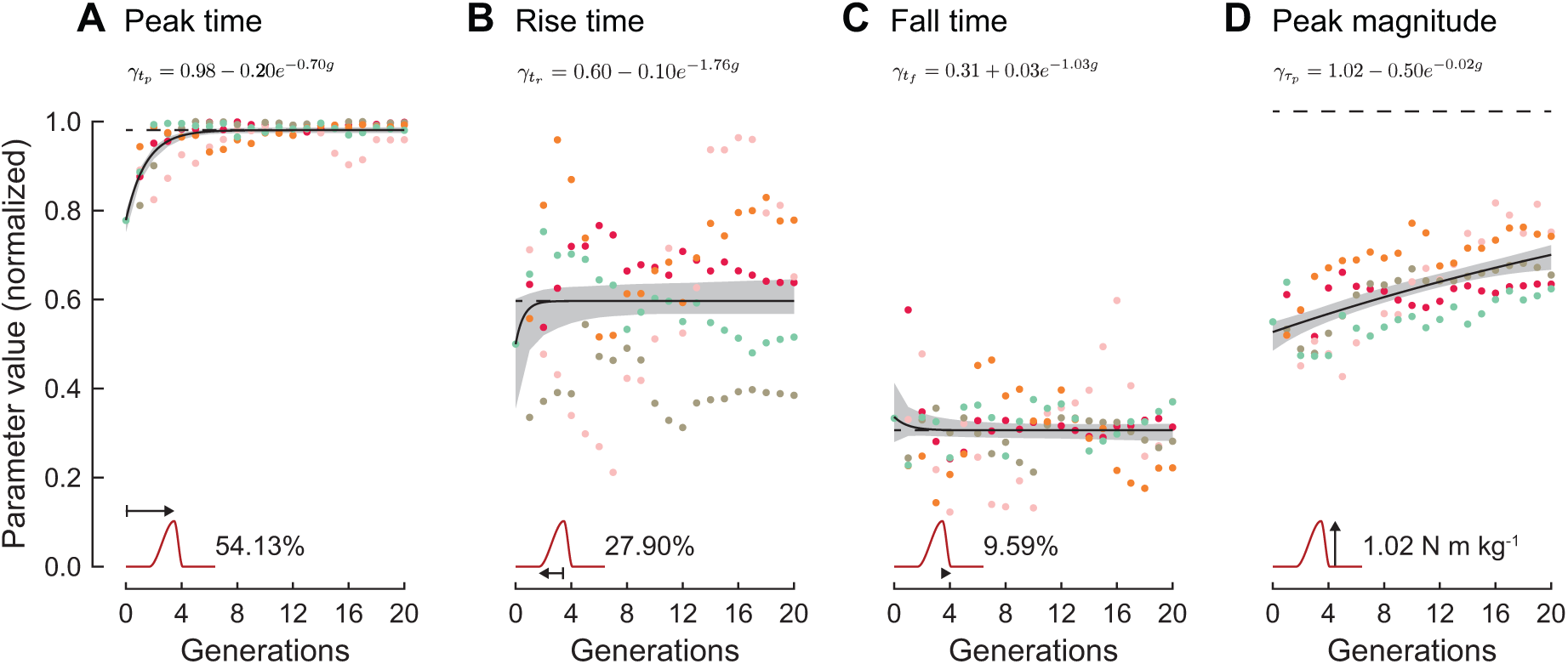
Optimized parameter values by generation for the *continued optimization* group and corresponding exponential fits. (*A*) Peak time, as a percentage of stride time, quickly converged to 54.1% of stride, with participants averaging 54.3 ± 0.7% after 20 generations (Mean ± S.D.). (*B*) Rise time appeared to be subject-specific, with the model converging to 27.90% of stride (average after 20 generations: 27.8 ± 4.5%). (*C*) Fall time also quickly converged to 9.59% of stride (average after 20 generations: 9.7 ± 0.9%). Peak time and fall time are coupled when peak time occurs late in stance, which may explain the stable fall time values. (*D*) Peak magnitude was predicted to converge at 1.02 N m kg^−1^, but participants optimized to lower values, 0.68 0.06 N m kg^−1^, after 20 generations of optimization. For all panels, individual participants are represented by distinct colors, and the exponential models, written at the top of each panel, are shown in black with corresponding 95% confidence bands in gray.

Based on the metabolic cost of optimal assistance, which encodes all of these factors as well as the time for the person to adapt to the exoskeletons, participants in the *continued optimization* group adapted to customized assistance in 330 minutes (Fig 8, IQR, 184 to 557 minutes). While the single exponential model suggests a longer training time compared to generic assistance, this estimate captures the effects of several phenomena of varying timescales, some of which have not yet converged by the end of the experiment. Although multiple timescales for motor adaptation have been demonstrated for other contexts (*33*), we hypothesize that convergence in metabolic cost for customized assistance in this case may reflect a decrease in variability in the tested parameters rather than true convergence of the human-exoskeleton system.

**Figure 8:**
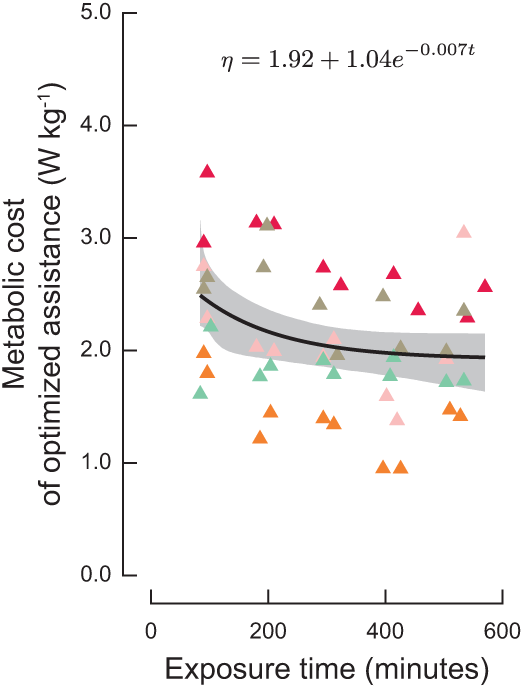
Metabolic cost of optimized assistance. The metabolic cost and corresponding exponential model, written at the top, for the optimized assistance condition for the *continued optimization* group. Individual participants are represented by distinct colors. The exponential model is shown in black with corresponding 95% confidence band in gray.

There is some evidence that the optimization algorithm could converge quickly for certain parameterizations. The timing parameters may have been easier to learn and thus faster to converge. The initial value for peak time was far from the final value, and starting with a better seed could speed optimization. The importance of this parameter may also explain why participants performed so well without the optimization fully converging in both our experiment and in other human-in-the-loop optimization studies (*24*). However, peak torque magnitude steadily increased throughout the experiment without converging. Because novices are often unable to accept high torques, simply seeding the optimization with a higher magnitude may not yield favorable results. Another experiment with fully trained participants could also be conducted to fully understand how quickly the magnitude of assistance can be optimized with human-in-the-loop optimization without the confounding effects of human motor adaptation.

### Optimized assistance had higher mechanical power than generic assistance

The optimized torque trajectories provided significantly more mechanical power than the generic profile (Fig 6B, repeated measures ANOVA, *P* = 2.9*e* − 10). While assistance profiles with higher device input resulted in lower energy output, mechanical power appeared to provide diminishing returns on metabolic cost reductions. The average reduction in metabolic cost for this group in response to generic assistance was 0.95 W·kg^−1^ while the reduction in metabolic cost in response to the optimized assistance was 1.21 W·kg^−1^, compared to a doubling in mechanical power, with 0.11 W·kg^−1^ for generic assistance and 0.23 W·kg^−1^.

We did not design this study to examine the complicated relationship between mechanical power and metabolic cost, but further research should be done to understand the complexities of energy transfer in the human-exoskeleton system. While we collected metabolic cost for candidate profiles with a wide variety of mechanical power, the optimization systematically adjusted parameters so as to reduce energy cost over time, leading to a sample of control laws that were necessarily correlated with lower metabolic cost. A full characterization of the relationship between metabolic cost, mechanical power, training, and other exoskeleton and user characteristics would make an important contribution to exoskeleton research and should be the subject of future studies. To fully explore the relationship between mechanical power and metabolic cost, exoskeleton parameters could be sampled over a large range of mechanical power and tested on expert exoskeleton walkers to determine how differing levels of mechanical assistance affect energy economy rather than selecting for conditions which reduce metabolic cost. For example, our results indicate that timing of assistance is an important factor, so varying the timing of the injected work may better explain the relationship between device power and metabolic energy consumption. Future studies on this subject should ensure that subjects are fully trained given the coupling of adaptation and mechanical power (Fig 5). Our lab has previously conducted experiments to characterize the relationship between device power and metabolic cost (*12*), but the participants were not fully trained and we now know that we can sample a larger space with expert exoskeleton users.

### Scarcity of exoskeletons, trained exoskeleton participants, and open-access exoskeleton data limit research

These results point to a need for fully-trained users for future exoskeleton studies. Experimental protocols should be longer to ensure users are fully adapted. For this study, with time for setup and breaks between trials, the average participant needed roughly 29 hours of lab time for 9.2 ± 1.2 hours of walking. These participants were healthy and active, and all were able to complete the 72-minute trials without breaks. Populations with decreased mobility, such as patient populations or elderly populations, would likely require increased time in the lab to complete the same protocol. Studying the long-term effects of exoskeleton assistance is also an important next step, but was outside the scope of this study.

Improving and speeding training is an important and relevant area for future research. While we identified some features of training that resulted in better, faster metabolic cost reductions, we cannot deem one of them an “optimal” strategy nor can we claim a universal adaptation for all assistive devices. A combination of strategies may be more effective, or an entirely different protocol based on step-to-step variations or a designed curriculum may yield even better results, dependent on the devices being used. We gave participants minimal instruction, but we know that targeted instruction and biofeedback can result in better outcomes in other training paradigms (*55, 65*). Analysis of the provided data in the Supplementary Materials may lend insights into which aspects of locomotion could be used as targets for biofeedback. Participants were also not allowed to consume media or other forms of distraction during this experiment, but increasing intellectual engagement or entertainment may make recruiting participants easier and may even result in faster adaptation.

Simulations of exoskeleton assistance could also facilitate experiments by identifying beneficial strategies before testing a population. Ensuring accuracy in simulations of human-exoskeleton systems is difficult now due to the lack of data on the response to exoskeletons. While we did not perform any biomechanics analyses in this study, we collected a complete biomechanics dataset (Supplementary Materials) so that future researchers can advance the field without the overhead of performing these lengthy experiments. This set is limited to a young, healthy population using bilateral ankle exoskeletons, and training populations with movement disorders or adapting to more complex devices may be an even slower process, so collecting and sharing these data sets would invite more research into each paradigm.

There are limitations to this study, primarily due to the supply of subjects. A larger sample size would have more power, and extending the protocol may have answered more questions such as how peak magnitude converges, but the difficulty in recruiting subjects would be increased. Because there is a small number of labs researching exoskeleton assistance and the overall costs to train participants is high, the future of exoskeleton research may rely on designing studies with fewer participants (*58*) or amassing and testing the same pool of expert exoskeleton users. With careful cultivation of a variety of participants or a better understanding of how individual differences affect the response to assistive devices, the results from such studies could be generalizable.

## Conclusion

This experiment highlighted the importance of training and adaptation on exoskeleton performance and illustrated the benefits of device customization. Participants who received moderate levels of variation during training significantly reduced their metabolic rate with customized assistance, of which half was due to training and one quarter to customization. The time required to stabilize metabolic cost was significantly longer than previous exoskeleton research and depended on the type of training experienced. These results demonstrate that people can learn how to exploit the exoskeleton provided they have proper training and sufficient time to explore. The human-exoskeleton system is complex, even without the confounding effects of time-varying motor adaptation. Giving participants more time to adapt to exoskeleton assistance should result in better and more reliable outcomes for assistive device studies. Some previous exoskeleton failures may have actually been failures of training.

## Materials and Methods

### Participants

Fifteen participants (5 female, age = 24.3 ± 3.2 yrs; body mass = 71.0 ± 11.4 kg; height = 1.72 ± 0.09 m) with no prior history of movement disorders completed this study. This study was approved by the Stanford Institutional Review Board and written consent was provided by all subjects before participation.

No participant had worn an exoskeleton prior to the study. Participants were randomly assigned to an adaptation group described below before the first data collection (Fig 1C). Sixteen participants were recruited. One participant ceased communication with the research staff and thus was dropped from the study. One participant ended the study after five days due to discomfort, and one participant ended after five days because they appeared to have converged. Both participants who completed five days of experiments were included in the analyses presented here. All other participants included in the study completed at least six days of experimentation.

### Exoskeleton hardware and control

Experiments were conducted using a bilateral ankle exoskeleton emulator (*27*). The exoskeleton end-effector consisted of a lightweight (0.88 kg) instrumented frame attached to a commercially-available running shoe (Fig 1A). The device contacted the participant at the foot and at the shank just below the knee. The exoskeletons were actuated by powerful off-board motors via a flexible Bowden cable transmission. The participant chose from men’s shoe sizes of 7, 9, or 11 and were given padding if the shoe was too large. The same model and size of shoes used in exoskeleton trials were used for normal walking trials.

During stance, a torque trajectory was prescribed by magnitude and time-based parameters (*24*) and was achieved with iterative learning low-level control (*66*). The trajectory was defined by peak time, the time at which the peak magnitude occurs as a percentage of the average stride time; rise time, the time for the trajectory to rise from 0 N·m·kg^−1^ to the peak magnitude as a percentage of the average stride time; fall time, the time to drop from the peak magnitude to 0 N·m·kg^−1^ as a percentage of stride time; and the peak magnitude in N·m·kg^−1^. The trajectory was smoothed by a cubic spline between the nodes. The control was triggered by a pressure sensor activated at heel strike. Iterative learning control exploits the periodicity of gait and has been used effectively with these devices (*8, 24*), with the root-mean-square error in torque tracking for this study as 2.5 N·m·kg^−1^, which is equivalent to 6.4% of the average peak torque in the generic assistance condition.

The swing phase, triggered by toe-off, was controlled as a function of ankle position. The ankle angle was tracked by a rotary encoder (Renishaw, Gloucestershire, UK) and used to determine toe-off. The motor was controlled to track the joint position with added slack to avoid interfering with the participant’s natural kinematics.

### Protocol

All participants completed a protocol shown in Fig 1C. Participants first completed a quiet standing trial on each day to determine their resting metabolic rate. On the pre-test day, participants completed a double-reversal validation test with the following trials: a normal walking trial with unaltered shoes to match the shoes in the exoskeleton, a zero torque trial with the exoskeleton worn but in a zero torque mode, and a generic assistance trial with a fixed generic torque profile. All walking trials were completed on a split-belt treadmill set at 1.25 m·s^−1^. The only verbal instructions were to “walk comfortably” and to “let the device do the work for you.”

On each subsequent day of the study, participants first completed a 72-minute adaptation trial with a 2-minute warm-up, based on random assignment, then a series of validation trials. Participants were given the opportunity for a break at each quarter of the adaptation trial, but were instructed that they could stop at any time. All groups completed the validation trials from the pre-test with other validation trials added for specific training groups. Each validation trial was six minutes in length. Participants were given a 5-minute break between trials.

The generic assistance profile was determined through pilot testing. The pilot experiment, described in greater detail in the Supplementary Materials, was a 2-day protocol with ten participants. Participants walked in bilateral ankle exoskeletons over one day of habituation and one day of human-in-the-loop optimization (*24*). The generic assistance profile for the experiment presented in this paper was defined by the average optimized parameters: peak time was 52.9% of stride, rise time was 26.2% of stride, fall time was 9.8% of stride, and, for participant mass *M* (kg), the peak torque magnitude was 0.54*M* N·m·kg^−1^.

#### Continued Optimization Training Group

Five participants (2 female, age = 23.4 ± 1.1 yrs; body mass = 70.2 ± 6.4 kg; height = 1.70 ± 0.11 m) were assigned to the *continued optimization* training group. During the adaptation trial on each training day, participants experienced four generations of human-in-the-loop optimization. The initial seed on the first day was set with peak time at 45% of stride, rise time at 25% of stride, fall time at 10% of stride, and peak torque magnitude at 0.55*M* N·m·kg^−1^, where *M* is the participant’s mass in kilograms. The optimization on each subsequent day continued from the optimized parameters from the previous day.

Each generation featured nine control laws, beginning with the generic assistance profile, followed by seven randomly sampled control laws, and ending with the mean of that generation. Each control law was experienced for two minutes, so each generation was 18 minutes in length. A sound signaled the beginning of a new control law, and the low-level torque tracking was reset to mitigate the effects of previous control laws. The generic assistance profile was tested at the beginning of each generation to track motor adaptation. To ensure comfort and mitigate the changes in control law, the seven randomly sampled control laws were sorted by peak torque magnitude.

At the end of each adaptation block, a new set of optimal parameters was generated and tested during the validation trials. The validation trials contained the trials from the pre-test, i.e., normal walking trials, zero torque trials, and trials in which the exoskeleton applied generic assistance. For this training group, participants also experienced the optimized profile from the end of that day’s adaptation block, as well as the optimized profile from the end of the first day of adaptation.

#### Static Training Group

Five participants (2 female, age = 23.6 ± 3.5 yrs; body mass = 67.6 ± 14.1 kg; height = 1.71 ± 0.08 m) completed the *static training*. Each control law during the adaptation block was the generic assistance profile, but with a sound to signal every two minutes. Because participants were blinded to their group assignment, they were unaware that all control laws were identical. The validation trials were strictly the trials from the pre-test, i.e., normal walking trials, zero torque trials, and trials in which the exoskeleton applied generic assistance.

#### Re-optimization Training Group

Five participants (1 female, age = 25.8 ± 4.1 yrs; body mass = 75.6 ± 13.0 kg; height = 1.76 ± 0.06 m) were assigned to the *re-optimization* training group. The adaptation trial was identical to the *continued optimization* training protocol, but the seed on each day was set to the same initial seed, i.e., peak time at 45% of stride, rise time at 25% of stride, fall time at 10% of stride, and peak torque magnitude at 0.55*M* N·m·kg^−1^, where *M* is the participant’s mass in kilograms. The control laws were randomly sampled so that participants experienced a new set of controllers from the same distribution on each day. The optimized parameters generated by the adaptation period were also tested in the validation trials. The validation trials were the same as for the *continued optimization* group, i.e., normal walking trials, zero torque trials, and trials in which the exoskeleton applied generic assistance, the optimized profile from the first day of optimization, as well as the optimized profile from that day of optimization.

### Data collection

Participants wore tethered bilateral ankle exoskeletons (Fig 1A) (*27*). The exoskeleton assistance was governed by a torque pattern parameterized by four parameters: peak magnitude, peak time, rise time, and fall time (*24*). The control loop ran at 1000 Hz on a real-time computer (Speedgoat, Liebefeld, Switzerland).

Standard biomechanics data including metabolic energy consumption were collected. Participants refrained from all food and drink except for water for at least three hours before they arrived for the experiment, which resulted in approximately four hours of fasting before the first trial. Volumetric oxygen consumption and carbon dioxide expulsion were measured throughout the experiment using an indirect calorimetry device (Cosmed Quark CPET, Rome, Italy) and streamed to the real-time exoskeleton controller. Metabolic energy consumption was computed using a standard equation (*67*). During optimization, the estimated metabolic cost of each controller was the result of a first-order dynamical model fit to single-breath data (*24, 68*). This method has been shown to result in low estimation errors, with an average of 4% error in a previous study (*24*). The optimization algorithm used these estimates to rank the control laws to minimize metabolic cost.

Net metabolic cost, computed offline after the experiment, was used in the analyses presented in this paper. A quiet standing trial was conducted at the beginning of the experiment after set-up to guarantee participants were at a fasted, rested state. Quiet standing was subtracted from all measurements of metabolic cost to yield the net metabolic cost of walking. Validation trials were six minutes in length, of which the second half, i.e., the final three minutes, were used for all analyses. The trial length and averaged measurement are standard practice in biomechanics and provide the truest representation of metabolic cost for steady state movement. The validation trials, which were two minutes in length, are represented by an estimate of steady-state metabolic rate based on a first-order approximation (*42, 68*). These estimates are used to determine optimal parameters during the experiment and have been shown to produce low (4%) error in predicting steady state metabolic cost (*24*).

Motion capture, electromyography, and force data were time-synced and recorded separately from the metabolic and device data. Motion capture data were recorded using 55 markers (Vicon, Oxford Metrics, Oxford, UK). Electromyography was recorded with Trigno wireless sensors on eight muscles on each leg (Delsys, Boston, MA, USA). Participants completed all walking experiments on an instrumented split-belt treadmill (Bertec, Columbus, OH, USA) which also provided ground reaction forces and moments. Ground reaction forces and moments were streamed to the computer running the real-time controller as well as recorded with the motion capture measurements. The biomechanical data were periodically recorded throughout adaptation and recorded for the entirety of the validation trials. These data were not included in this paper due to space constraints, but links to the data can be found in the Supplementary Materials.

### Statistical analysis

All statistical analyses were performed after the experiment in R 3.6.2 (*69–72*).

To determine the effects of training and training type on the metabolic cost of assistance, a 3-way ANOVA, followed by a stratified analysis on the post-adaptation data, was performed (Fig 2). For the 3-way ANOVA, the factors were day (pre-test, post-adaptation), training group (*continued optimization*, *static training*, *re-optimization*), and condition (normal shoes, zero torque, generic assistance, and optimized assistance). All three factors, i.e., day, group, and condition, were significant, indicating that training, training type, and assistance levels resulted in different levels of energy cost. To determine the effects of training group on condition, we did a subanalysis on the post-adapted states followed by the Holm-Šidák step-down correction for multiple comparisons, similar to an unpaired *t*-test with adjusted *P*-values for multiple comparisons. The post-adapted states were the validation trials from the final two days of the experiment, which occurred after the *continued optimization* group had reached expertise with optimized assistance. To isolate the effects of assistance and training on metabolic cost within training groups, paired *t*-tests followed by the Holm-Šidák step-down correction were performed. For all analyses, the significance level was *α* = 0.05.

Analyses of the optimized control laws for the *continued optimization* group are presented in the main text, and analyses for the *re-optimization* group can be found in the Supplementary Materials. A one-way repeated measures ANOVA was computed to determine differences in net exoskeleton power between conditions (Fig 6). The post-adapted states were the validation trials from the final two days of the experiment, as described for the metabolic cost analyses. A paired *t*-test was computed to determine differences in parameter values after twenty generations for the *continued optimization* group compared to generic assistance.

Exponential models were used to characterize human motor adaptation (Figs 3, 5, 8) and evolution of the optimal parameters (Fig 7). Changes due to adaptation were characterized by a single exponential curve of the form *η* = *a* + *a*_0_*e^−ωt^*, where *t* represents time in minutes, *a* is the steady-state response, *a*_0_ is a scaling factor, and *ω* is the time constant. The outcomes of interest, represented by *η*, were metabolic cost (Figs 3, 8) for generic and optimized assistance, respectively, and net exoskeleton power (Fig 5) for generic assistance conditions during validation and during the adaptation trial. The same model was used for the evolution of the parameters (Fig 7), where *η* was the value of the parameter normalized to its respective range and *t* was the number of generations. The model parameters {*a*, *a*_0_, *ω*} were estimated using a weighted nonlinear least squares algorithm (*70*). Validation trials were weighted three times the adaptation trials for these models because validation trial estimates were computed over a duration triple the length of the adaptation trials. Model significance was determined by an ANOVA model comparison with the constant model *η* = *a*, which is standard practice for linear and nonlinear regressions. For all model comparisons, the significance level was *α* = 0.05.

Bootstrapping was used to determine confidence intervals on the exponential models and to characterize the adaptation rates (*71, 73*). For a given model determined by nonlinear least squares, the residuals were sampled and replaced; these residuals were added to the model estimates to determine a new data set, which was then used to compute a new exponential model. This process was completed up to 10000 times, depending on how often the model converged. These models were then used to determine the point-wise 95% confidence interval for the exponential model. This bootstrapping technique has been used in other biomechanical studies and has been shown to better capture the randomness in the underlying model (*73, 74*). The learning rates were computed as the time in minutes for the model to reach a value within 5% of the predicted asymptote. The learning rates presented in this paper were computed for the model determined by nonlinear least squares, and the corresponding confidence intervals were determined by the bootstrapped replicates (Fig 4). Because the bootstrapped learning rates did not follow a normal distribution, according to the Shapiro-Wilk test (*P <* 2.2*e* − 16), we performed the Kruskal-Wallis tests, followed by Dunn’s tests with *α* = 0.05 (*72*), to determine differences across training groups.

## Acknowledgments

We thank members of the Stanford University Biomechatronics Laboratory, in particular A.S. Voloshina, for their patience and assistance in data collections and insights on the work, as well as the Stanford Department of Statistics Consulting services for their guidance on the statistical methods.

## Author contributions

S.H.C. and K.L.P. were responsible for the design of this experiment. K.L.P. conducted the experiments and analyzed the data. K.L.P. and S.H.C. prepared the manuscript.

## Data and materials availability

All data are available in the paper or in links found in the Supplementary Materials.

