## Supplementary Materials for "How adaptation, training, and customization contribute to benefits from exoskeleton assistance"

This pdf file includes:

- Methods and Results
- Figs. S1 to S9
- Table S1
- Captions for Movies S1 and S2
- Links and captions for Data S1 to S15
- References

### 1 Supplementary Methods

#### 1.1 Pilot test to determine generic assistance profile

Ten participants with no prior exoskeleton experience were recruited for the two-day protocol. Participants wore bilateral ankle exoskeletons as in the main study.

##### 1.1.1 Protocol

The first day of the experiment began with a familiarization period, followed by six validation trials. The familiarization trial was ten minutes in length, with each minute varying one of the parameters: peak time ($\%$ stride), rise time ($\%$ stride), fall time ($\%$ stride), and peak magnitude (N·m·kg$^{-1}$). Some parameter changes were done slowly, to allow for a comfortable transition, or instantaneously, to familiarize participants with the human-in-the-loop optimization protocol. The speed at which parameters were increased during the ramp phases was dictated by the experimenter and their assessment of participant comfort.

The validation trials included baseline trials of normal shoes and the zero torque mode, as well as a generic profile. This profile was the average of the parameters from a unilateral human-in-the-loop optimization protocol (*24*), with a peak time of $50.4\%$ of stride, a rise time of $23.3\%$ of stride, a fall time of $12.3\%$ of stride, and a peak magnitude of $0.74M$ where $M$ is the participant's mass in kilograms.

On the second day, the participants underwent the optimization trial followed by a series of validation trials. The optimization was 64 minutes, with four generations of eight randomly sampled control laws, experienced for two minutes (*24*). Validation followed the optimization trial, with the baseline trials, the average torque described above, and the optimized control law.

##### 1.1.2 Results

Participants initially struggled to walk with assistance after the familiarization and exhibited no reduction in energy cost with assistance on the first day (Fig. S1, paired $t$-test, $P = 0.94$). After four generations of optimization in this one-day protocol, participants reduced their energy cost with both the generic assistance and optimized condition by $18.2\%$ and $18.6\%$ compared to the zero torque condition, respectively. All participants optimized to profiles characterized by a peak that was lower in magnitude (paired $t$-test, $P = 3.9e-4$) and later in time (paired $t$-test, $P = 0.006$) than the generic profile based on the unilateral experiment. The average profile determined by these parameters was defined by a peak time of $52.9\%$ of stride, a rise time of $26.2\%$ of stride, a fall time of $9.8\%$ of stride, and a magnitude of $0.54M$ where $M$ is the participant's mass in kilograms. These parameters differ from those determined by the unilateral exoskeleton study (*24*). Because the original, published human-in-the-loop optimization study was conducted with a unilateral ankle exoskeleton (*24*), we used the average profile determined by the pilot bilateral exoskeleton experiment.

#### 1.2 Subject characterization tests

We were interested in discovering participant-specific qualities that could predict some measure of exoskeleton performance. We considered biometric information (body mass in kilograms), experimental condition (training group), grit (*75*), and a measure of dexterity (*76*). The outcomes we sought to predict for the *continued optimization* and *static training* groups were the metabolic cost of generic assistance during the pre-test (W·kg$^{-1}$), the metabolic cost of generic assistance after adaptation (W·kg$^{-1}$), and the individual adaptation time for generic assistance (Fig. S5, minutes). For the *continued optimization* group, we included the metabolic cost of optimized assistance (W·kg$^{-1}$) and the optimized peak magnitude value (N·m·kg$^{-1}$). We did not conduct these analyses for the *re-optimization* group because we do not believe they adapted in the same way.

The independent variables were randomly assigned (experimental condition), self-reported (body mass, grit), or measured in a separate test (dexterity). Training group was determined before the participant entered the lab, and body mass was given by the participant upon arrival. Grit, a measure of perseverance and resilience, was determined by a 12-question survey (*75*) given to participants after consent was given before the pre-test. Lyle, et al., developed a protocol and experimental apparatus to determine dexterity (*76*). Participants were instructed to push down on a wooden board attached to a spring that was prone to buckling for 16 seconds in 25 trials with breaks in between; we modified the apparatus by removing some of the restraints, which allowed for more freedom of movement during the test. The average reaction force (N) produced over the last five trials were used as the measure of dexterity. Because this test was added to the protocol after the experiment had finished for some participants, there is a chance that their dexterity may have changed in the time between the end of the main experiment and this additional test.

Multivariate linear mixed-effects models were computed for each outcome; training group was not included as an independent variable for optimized metabolic cost or peak magnitude. There were no significant ($P < 0.05$) models reported for any of the outcomes, and only the intercept was a significant factor for the metabolic cost of generic assistance during the pre-test ($P = 0.02$) and after adaptation ($P = 0.007$) and for the expertise time ($P = 0.047$), indicating that these outcomes had non-zero mean. These results show that the individual response to exoskeleton assistance cannot be described by these external factors. Athletic ability was also considered as a predictor but was not tested; in our experience, qualitative athletic ability is not a good predictor but a more rigorously-defined metric might relate to outcomes relevant to exoskeleton use.

#### 1.3 Exponential model analysis

There are two choices to be made in order to describe how people learn to use a device. The first is identifying the data to represent "motor learning" or "expertise," and the second is how to model that data. As described in the main text, motor learning, represented by averaged metabolic cost, was modeled as an exponential function of exposure time,

$$\eta = a + a_0 e^{-bt}, \quad (1)$$

computed by weighted nonlinear least squares with the starting point $a = 2, a_0 = 1, b = 0.02$.

Models of motor learning vary in form, but share some common characteristics. Learning is cumulative, so the representative curves are monotonic. Learning is also typically nonlinear, with rapid progress initially and small improvements at later stages. This can result in asymptotic behavior, which suggests that there is some final expert state. Several curves have been proposed, including an exponential function (*32*), a power curve (*19, 30*), and a double exponential function separating slow and fast learning processes (*34*).

We modeled study-level generic assistance data as a horizontal line, a line with nonzero slope, a power function, a single-term exponential function, and a two-term exponential function (Fig. S3). The linear models were computed using weighted ordinary least squares, and the nonlinear models using nonlinear least squares, where the six-minute validation trials were weighted three times as much as the two-minute adaptation trials (*70*). For the following models, $\eta$ represents the metabolic cost in W·kg$^{-1}$ and $t$ represents exposure time in minutes. As expected, the constant line

$$\eta = 2.53 \quad (2)$$

fit the data poorly (weighted residual standard error = 0.75), as did the line with nonzero slope

$$\eta = 2.78 - 0.001t \quad (3)$$

(weighted residual standard error = 0.72). For the power function, to prevent division by zero errors, we transformed the independent variable $T = t + 6$. The power function

$$\eta = 1.90 + 2.05T^{-0.24} \quad (4)$$

and the single-term exponential function

$$\eta = 2.39 + 0.72e^{-0.017t} \quad (5)$$

both fit the data with low residual error (weighted residual standard error = $0.69, 0.69$, respectively). The two-term exponential function

$$\eta = 2.37 + 0.41e^{-0.008t} + 0.41e^{-0.050t} \quad (6)$$

did not improve the model fit (weighted residual standard error = 0.69), and therefore the additional parameters did not seem justified. The single-term exponential had the lowest weighted residual standard error of the models we tested, but this analysis demonstrates that other common motor learning models could be used to characterize adaptation to exoskeletons.

Metabolic cost of generic assistance, normalized to body mass, was used as a proxy for motor adaptation. The torque profile was held constant across participants to remove variations in assistance profiles as a confounding variable. Metabolic cost was normalized to body mass rather than the average metabolic cost of the zero torque conditions on that day, which was done for aggregate results in the main text. To determine if there was adaptation to the zero torque condition, a similar exponential analysis was performed, demonstrating that participants learned how to use the zero torque condition in 27 minutes (Fig. S2A). The zero torque condition is akin to wearing a large boot, so it is unsurprising that participants can quickly adapt to it.

Averaged metabolic cost was chosen because it was granular enough to capture the motor adaptation over minutes without obscuring any dynamics within the trial. While steady exponential improvements in energy cost have been demonstrated for exoskeleton walking over long trials using single-breath data (*17*), the dynamics of metabolic cost within a single trial in our study were dominated by the transient changes in metabolic cost associated with accelerating from standing to steady-state walking. We modeled metabolic cost of individual breaths as a function of exposure time using only the validation trials. This was to better compare between the other methods presented here, and model analysis of the validation using the method in the main text showed similar properties. The single-breath data predicted a steady-state metabolic cost of 2.38 $W{\cdot}kg^{-1}$, achieved in 122 minutes. These single-breath values were averaged for each trial and were therefore incorporated into the exponential model in the main text, so it is to be expected that an exponential model fit to single-breath data would yield similar results.

Metabolic cost was modeled as a function of exposure time. While developmental research indicates that exposure time is a better predictor of walking ability (*41*), exoskeleton research often uses "training days" as the independent variable (*18*). Adaptation was seen within a single day, which was one reason to avoid using whole days as the independent variable. We did however fit exponential models to the averaged metabolic cost of the generic assistance validation trials as a function of both testing day and calendar day. As a function of experiment day, with 0 for the pre-test day, participants were predicted to reach a steady-state metabolic cost of 2.38 $W{\cdot}kg^{-1}$ within 1.55 experiment days. Exoskeleton exposure by the end of the first training day would have been between 96 and 120 minutes, depending on the training protocol, so these models show a similar adaptation time and similar estimated steady-state cost (compared to 2.39 $W{\cdot}kg^{-1}$). Metabolic cost as a function of calendar time was predicted to achieve a similar steady-state cost (2.43 $W{\cdot}kg^{-1}$) in 2.29 days. Participants were required to have at least one rest day between experiment days, so the second day of adaptation would have occurred on day four at the earliest. For groups that experienced 120 minutes of exoskeleton exposure by the end of the first day of adaptation, this estimate of 2.29 days would indicate that some learning occurred after the experiment. While we know that consolidation can aid motor learning (*39, 77*), this experiment was not designed to characterize the effects of sleep on exoskeleton training. For this experiment, we were not able to constrain participants to a strict schedule, so analysis of calendar time provides little additional insight.

### 2 Supplementary Results and Discussion

#### 2.1 *Re-optimization* group analyses

In addition to treating the *re-optimization* protocol as a separate training curriculum, we were interested in the implications of reseeding and rerunning human-in-the-loop optimization protocols on an individual. Do participants reliably converge to the same parameters after four generations? If not, how do the optimized profiles change over time and what similarities exist in those optimized profiles? Because the metabolic cost of generic assistance was relatively unchanged throughout the experiment for this group, day-to-day variation within each participant represents variation in how the parameters are optimized on that day rather than the multi-day adaptation seen in the other groups.

Participants did not reliably converge to the same assistance profile after four generations (Fig. S8), and there were no discernible patterns in the resulting assistance profiles throughout the experiment. In general, participants converged to a late peak time of $53.4 \pm 11.3\%$ of stride. The spread in the optimized peak time was small, with participants only using a small percentage of the available space, i.e., the optimized peak time only ranged from $49.6\%$ to $54.8\%$ of stride. The other parameters were more varied. Average rise time was $24.7 \pm 15.2\%$ of stride, and average fall time was $10.0 \pm 7.3\%$ of stride; both similar to the initial seeds of 25% and 10% of stride, respectively, although the values varied across days. The average peak magnitude, $0.59 \pm 0.12$ $N{\cdot}m{\cdot}kg^{-1}$, was lower than the peak magnitude seen for the *continued optimization* group by the end of optimization, but higher than the average peak magnitude after four generations for that group ($0.55 \pm 0.09$ $N{\cdot}m{\cdot}kg^{-1}$). The mechanical power of the optimized conditions was also varied across participants ($0.22 \pm 0.12$ $W{\cdot}kg^{-1}$).

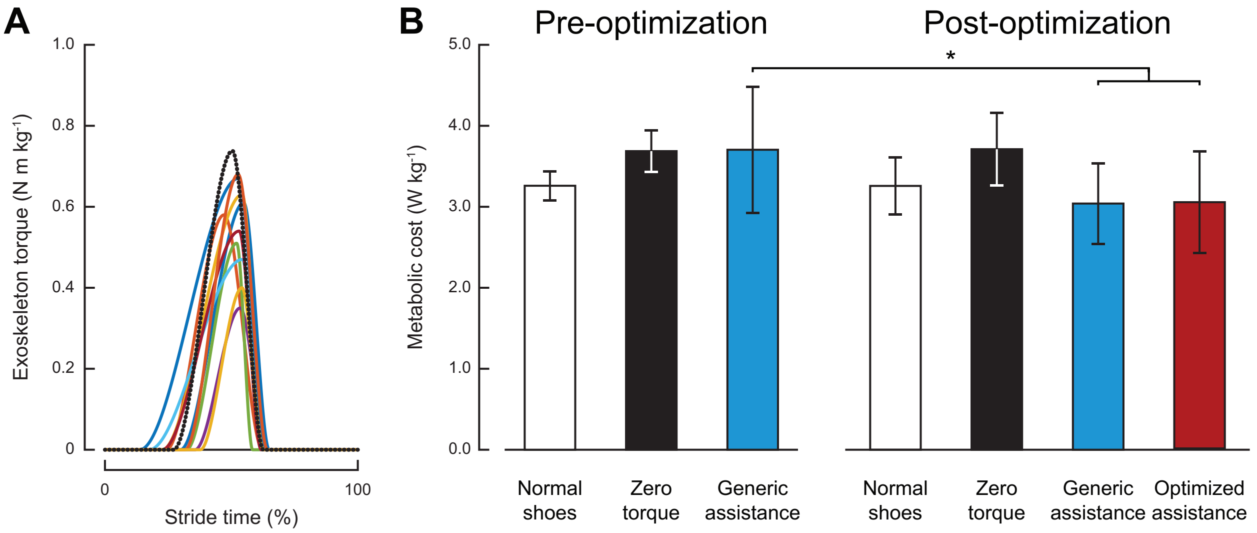


Figure S1: **Results from the pilot experiment to determine the generic assistance profile.** (*A*) Optimized torque profiles for the ten participants, in solid colors, and the unilateral profile used as the generic assistance condition (*24*), in a dotted black line. The peak magnitude of the optimized profiles was significantly lower than the peak magnitude of the unilateral profile (paired $t$-test, $P \ll 0.001$). (*B*) Metabolic energy cost measured during the pilot test. After the familiarization trial on the first day, there was a $0.3\%$ increase in metabolic cost with the generic assistance. By the second day of testing there was a $18.2\%$ reduction in metabolic cost with the generic assistance profile and a $18.6\%$ reduction in metabolic cost with the optimized profile.

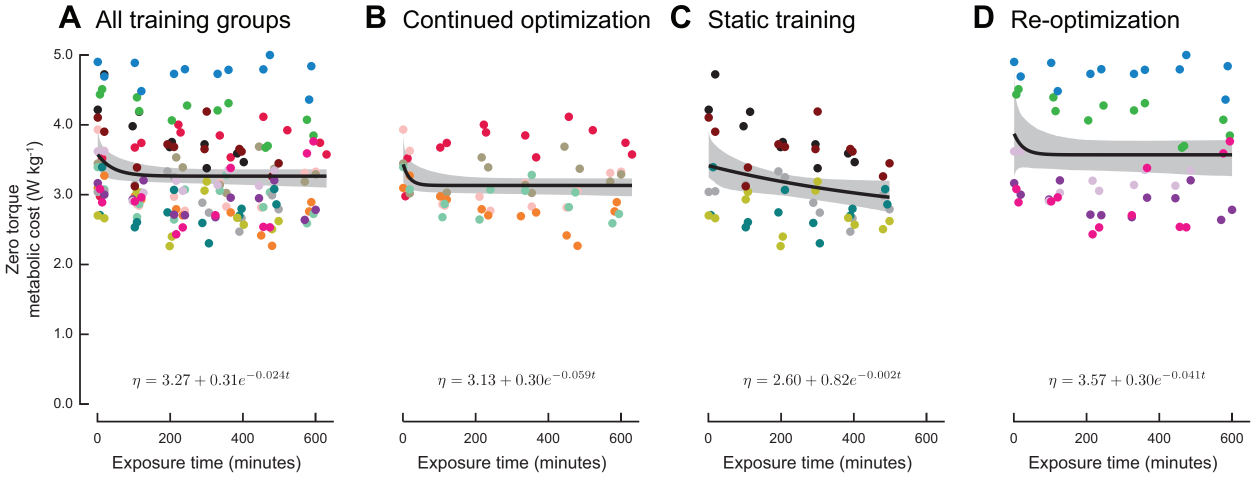


Figure S2: **Metabolic cost of zero torque.** Metabolic energy cost and corresponding exponential models of the zero torque conditions at (*A*) the study level and at training group levels for the (*B*) *continued optimization*, (*C*) *static training*, and (*D*) *re-optimization* training groups. The zero torque condition applies minimal force, so adaptation to zero torque is similar to adding mass and a rigid constraint on the end-effector rather than changing the joint kinetics. At the study level, participants achieve expertise within a half hour. The wide confidence intervals for the *re-optimization* and *static training* groups illustrate the subject-specific nature of zero torque adaptation; in the case of the *static training* group, this variability in response results in a model that predicts a longer adaptation time than what would be expected for the individual participants. For all plots, individual participants are represented by distinct colors. The exponential models, written at the bottom of each plot, are shown in black with corresponding 95% confidence bands in gray.

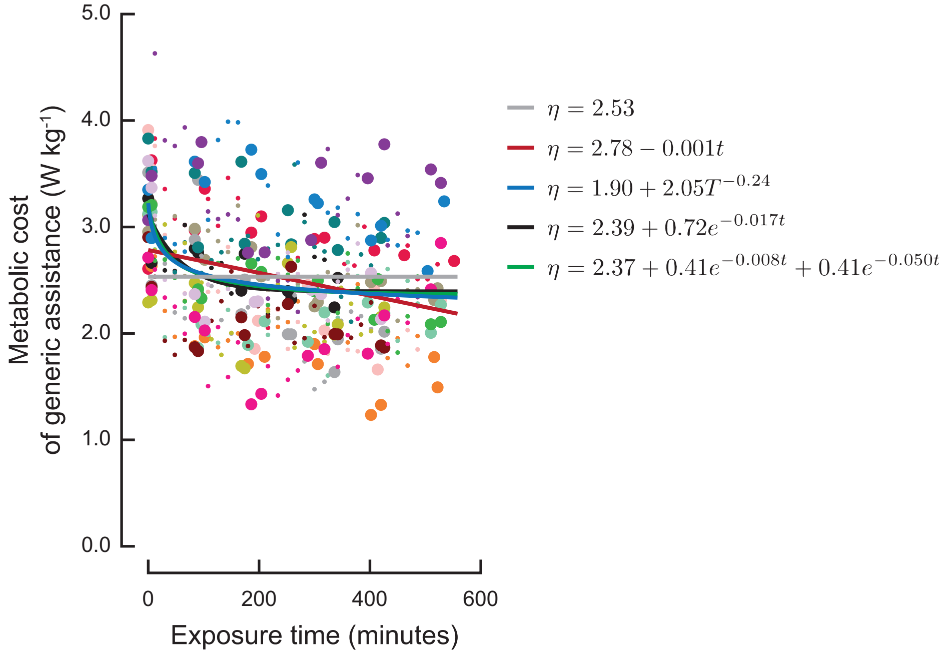


Figure S3: **Study-level model comparison to characterize the metabolic cost of walking with generic assistance.** Beyond the single-term exponential model described in the main text and shown here, a linear model, a two-term exponential model, and a power model were fit to the data; a baseline constant was added for reference.

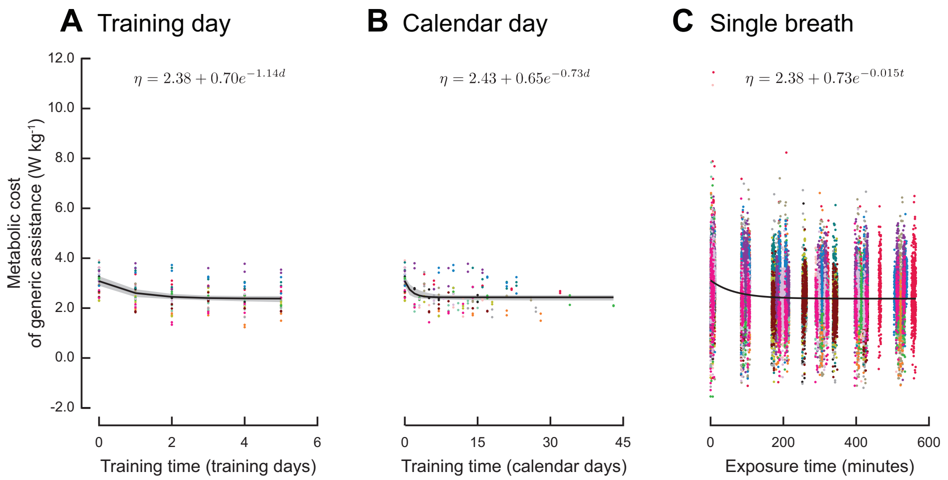


Figure S4: **Metabolic cost of generic assistance using differing measures of time.** Metabolic energy cost and corresponding exponential models of fixed generic assistance for (*A*) averaged metabolic cost as a function of training day, (*B*) averaged metabolic cost as a function of calendar day, and (*C*) single-breath metabolic power as a function of exposure time. The model in the main text predicted expertise at 109 minutes, which would have occurred early in the second adaptation period. Metabolic cost as a function of experiment day shows a similar adaptation rate, with participants reaching expertise at 1.55 days, where the pre-test day was considered day 0. Metabolic cost as a function of calendar day suggested an earlier convergence time of 2.29 days; because participants were required to rest for at least 24 hours between testing, we would expect this model to predict expertise between four and five days. The predicted time to expertise with the single-breath data of 122 minutes is similar to the model using the averaged data, but the noise in the measurements obscures the metabolic cost of each trial. Individual participants are represented by distinct colors, and only the 6-minute validation trials were used. The exponential models, written at the top of each plot, are shown in black with corresponding 95% confidence bands in gray; the confidence band for this model was very tight and so may be difficult to see.

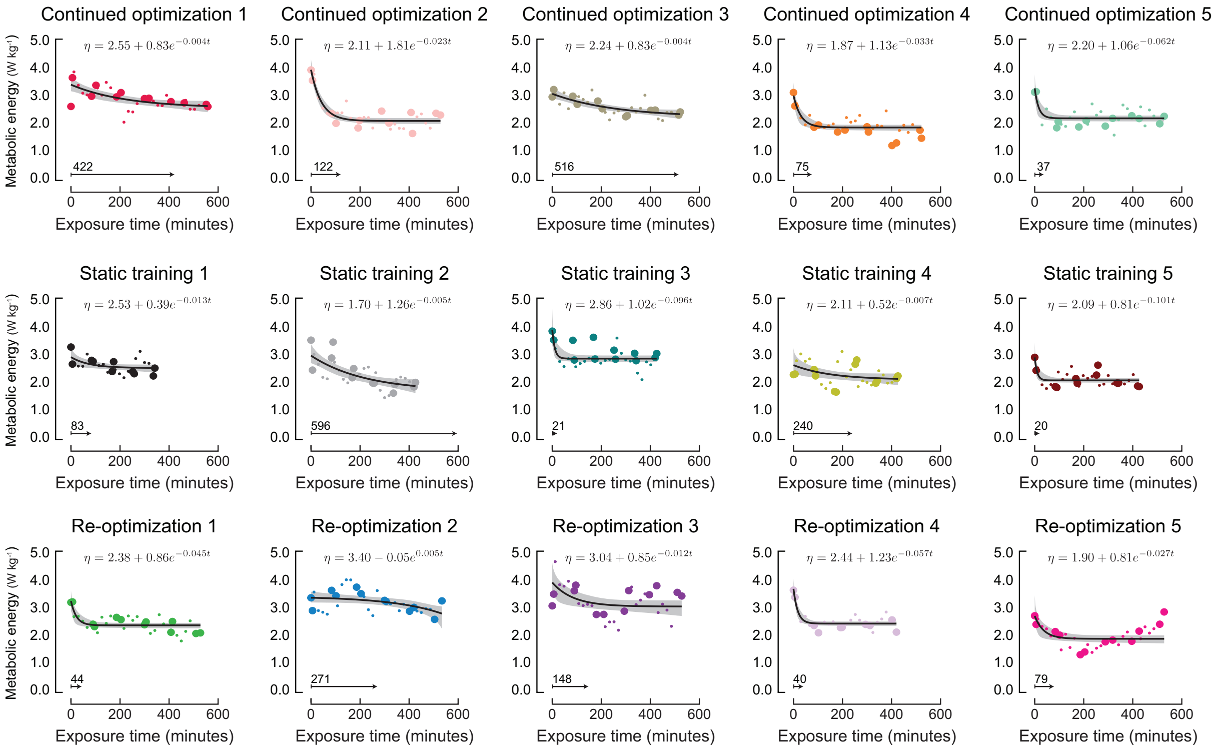


Figure S5: **Metabolic cost and corresponding exponential models of the fixed generic assistance conditions for individual participants.** Participants exhibit varied responses and therefore varying amounts of time to gain expertise. Most participants reduce their metabolic cost in an exponential decay throughout the experiment, reaching a steady-state value before the end of the experiment; one participant (the second *re-optimization* participant) is predicted to reduce metabolic cost without bound, which we do not expect to be the case if we were to continue the experiment. The 6-minute validation trials are represented by larger points than the 2-minute estimates taken during the adaptation trials. The exponential models, written at the top of each plot, are shown in black with corresponding 95% confidence bands in gray. The expertise time, or the time for the model to cross a threshold of 5% of the asymptote, is shown at the bottom of each plot.

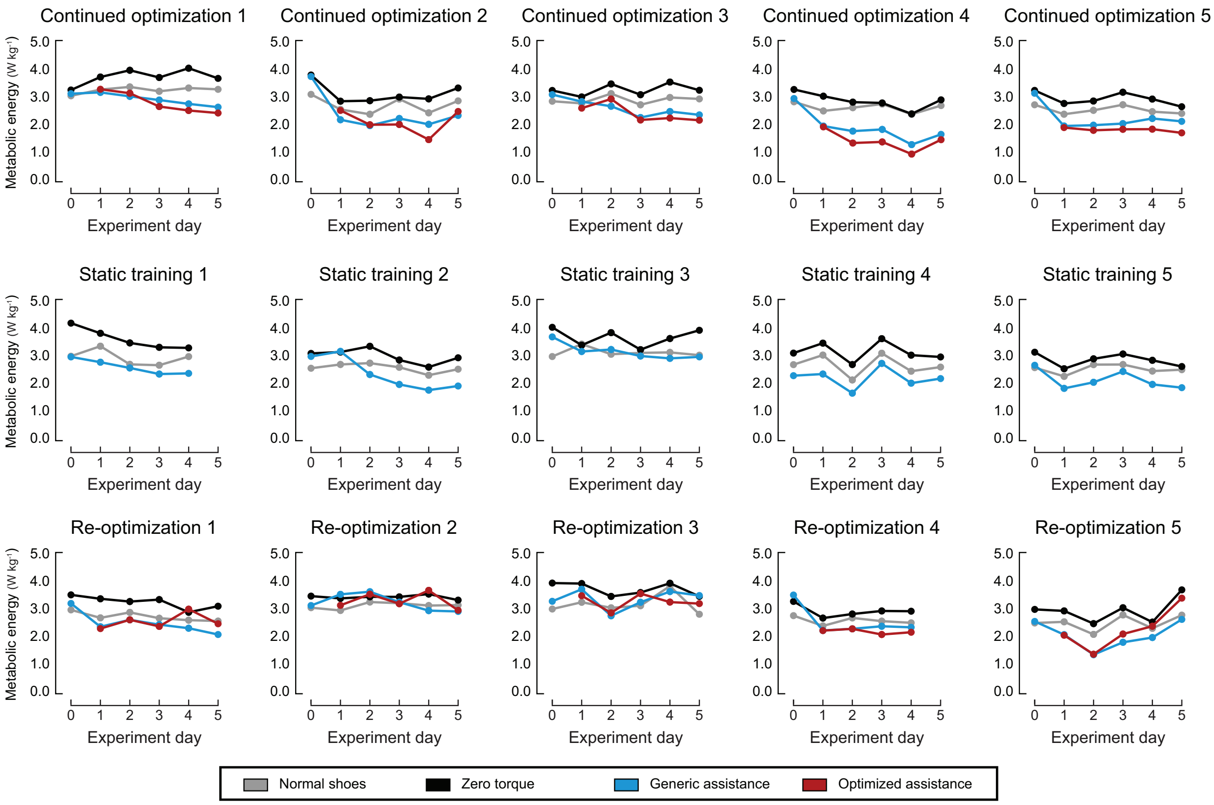


Figure S6: **Net metabolic cost of all conditions for each participant.** There are normal fluctuations in metabolic cost across days, as demonstrated by the cost of walking in normal shoes shown in gray. Much of the variation in the zero torque conditions tracks the day-to-day fluctuations seen in walking with normal shoes, especially toward the end of the experiment after participants have adapted to the zero torque condition.

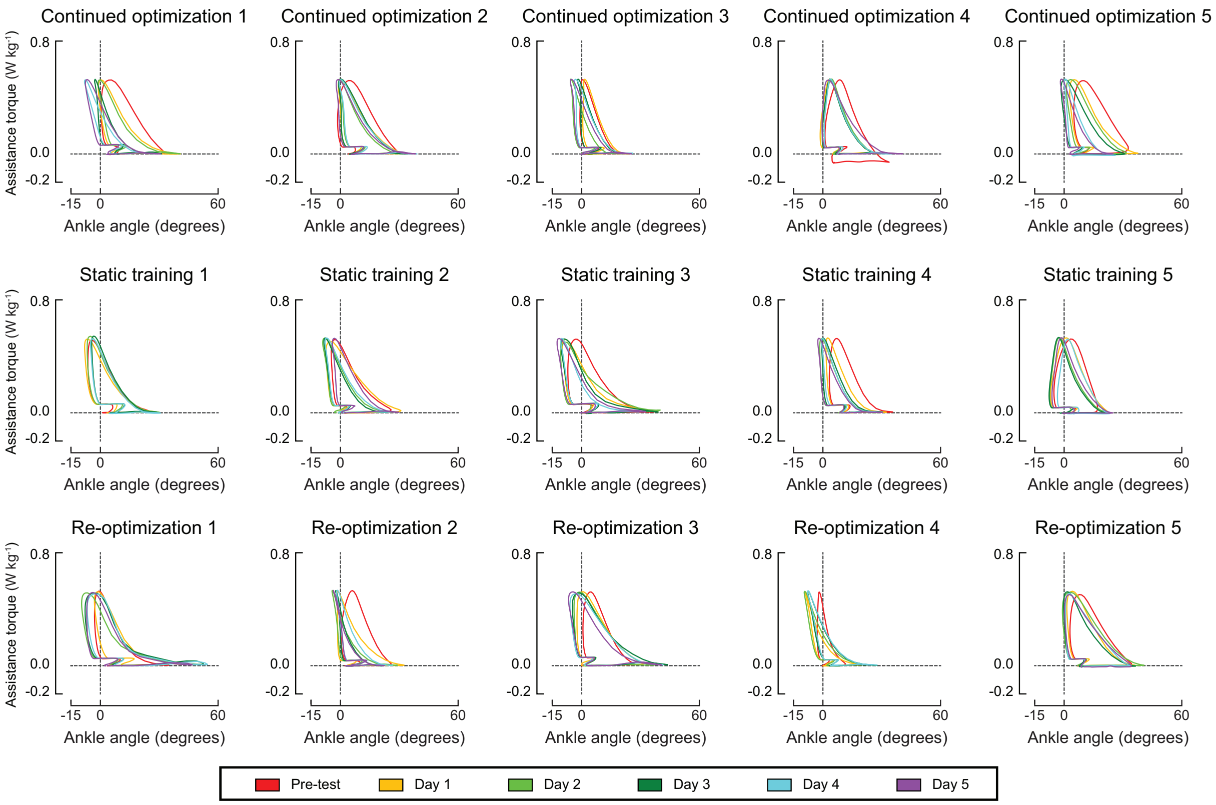


Figure S7: **Exoskeleton torque vs. ankle angle for generic assistance.** Each line represents the average trajectory across generic assistance validation trials for each day of the experiment. The area within the curve is the mechanical work from the device; work normalized by stride time is shown in Figure 5 in the main text. As stated in the main text, mechanical power decreased with training time at the study level and the group levels. Because the generic assistance torque profile is fixed, this reduction in mechanical power is the result of adaptations that affect ankle kinematics. Particularly evident when comparing the pre-test (red) to validation sessions from later training days (yellow through purple) is the larger ankle velocity during periods of torque application, which manifests as a wider work loop and a more plantarflexed angle at the instant of peak torque.

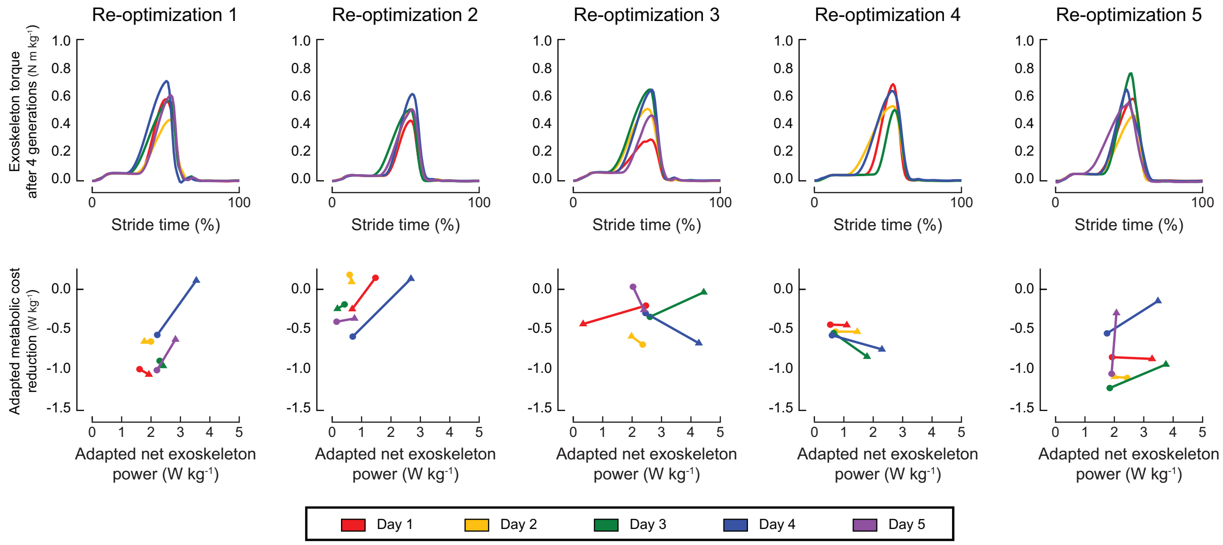


Figure S8: **Optimized and generic assistance profiles for each participant in the *re-optimization* group.** Profiles were considered "optimized" after the four generations of optimization on each day. On the top row, optimized torque trajectories ($N{\cdot}m{\cdot}kg^{-1}$) are shown in solid lines for each participant. The generic assistance torque trajectory is shown as a dotted line in black. On the bottom row, metabolic cost reduction is plotted against mechanical work for the generic assistance profiles, denoted ○, and for the optimal assistance profile, denoted △, on each day of testing. There were no obvious day-to-day trends across participants.

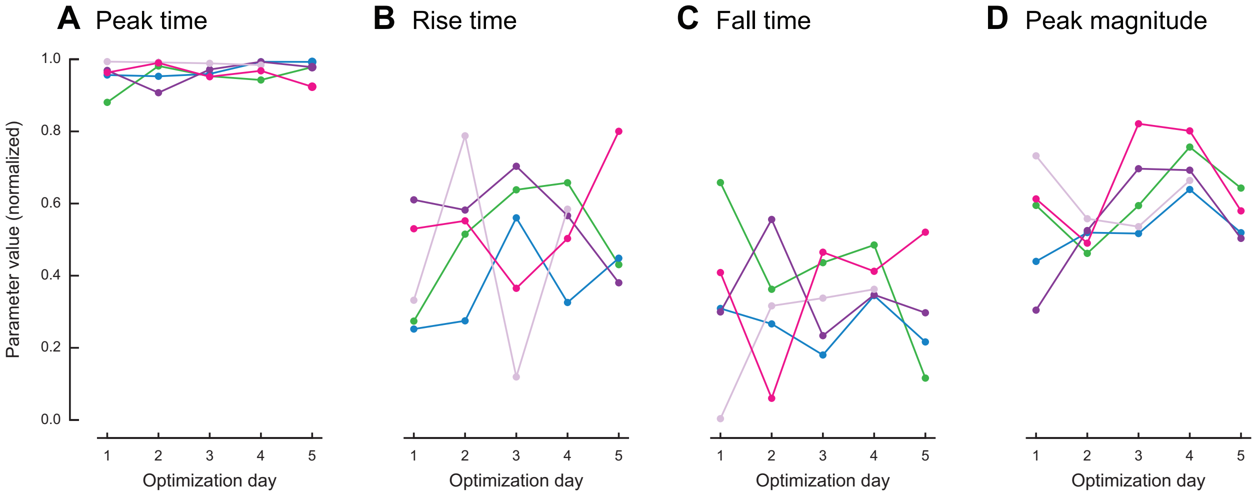


Figure S9: **Optimized parameter values at the end of each day of optimization for the *re-optimization* training group.** (*A*) Peak time, as a percentage of stride time, reliably converged to the end of stance. (*B*) Rise time and (*C*) fall time did not show any particular trends across day, nor within individual participants. (*D*) Peak magnitude also did not show a trend across days, but most participants reached their highest magnitude by the third or fourth day before reducing the magnitude on the final day of the experiment. For all panels, individual participants are represented by distinct colors.

Table S1: Subject Demographics

| Participant | Sex | Body mass (kg) | Height (m) | Age (yrs) | Grit |
|---|---|---|---|---|---|
| | | *Continued Optimization* | | | |
| 1 | F | 54 | 1.54 | 23 | 2.83 |
| 2 | M | 67 | 1.82 | 23 | 4.33 |
| 3 | F | 75 | 1.63 | 25 | 2.75 |
| 4 | M | 74 | 1.78 | 24 | 3.33 |
| 5 | M | 72 | 1.73 | 22 | 3.75 |
| **Mean ± SD** | 2 F, 3 M | 68.4 ± 8.6 | 1.70 ± 0.11 | 23.4 ± 1.1 | 3.40 ± 0.66 |
| | | *Static Training* | | | |
| 1 | F | 58 | 1.68 | 21 | 3.75 |
| 2 | M | 66 | 1.77 | 24 | 3.67 |
| 3 | F | 57 | 1.65 | 24 | 4.00 |
| 4 | M | 64 | 1.64 | 29 | 3.75 |
| 5 | M | 92 | 1.83 | 20 | 4.33 |
| **Mean ± SD** | 2 F, 3 M | 67.5 ± 14.4 | 1.71 ± 0.07 | 23.6 ± 3.5 | 3.90 ± 0.27 |
| | | *Re-optimization* | | | |
| 1 | F | 69 | 1.71 | 26 | 3.50 |
| 2 | M | 93 | 1.85 | 27 | 2.50 |
| 3 | M | 59 | 1.70 | 30 | 3.17 |
| 4 | M | 82 | 1.77 | 19 | 3.92 |
| 5 | M | 72 | 1.77 | 27 | 3.42 |
| **Mean ± SD** | 1 F, 4 M | 74.8 ± 13.3 | 1.76 ± 0.06 | 25.8 ± 4.1 | 3.50 ± 0.27 |
| **Total Mean ± SD** | 5 F, 10 M | 70.2 ± 11.9 | 1.72 ± 0.09 | 24.3 ± 3.2 | 3.60 ± 0.47 |

#### Movie S1

**Validation.** Validation trials for an example *continued optimization* participant.

#### Movie S2

**Adaptation.** Adaptation trial for an example *re-optimization* participant; the trial is similar to that experienced by the *continued optimization* group.

#### Dataset S1: purl.stanford.edu/jj710vy7867

Contains the device and metabolic data for the first *Continued Optimization* participant.

#### Dataset S2: purl.stanford.edu/ww452xb7000

Contains the device and metabolic data for the second *Continued Optimization* participant.

#### Dataset S3: purl.stanford.edu/mm626wf3265

Contains the device and metabolic data for the third *Continued Optimization* participant.

#### Dataset S4: purl.stanford.edu/zr858qp8088

Contains the device and metabolic data for the fourth *Continued Optimization* participant.

#### Dataset S5: purl.stanford.edu/hs191pw6736

Contains the device and metabolic data for the fifth *Continued Optimization* participant.

#### Dataset S6: purl.stanford.edu/st957pf8319

Contains the device and metabolic data for the first *Static Training* participant.

#### Dataset S7: purl.stanford.edu/mw935fz1170

Contains the device and metabolic data for the second *Static Training* participant.

#### Dataset S8: purl.stanford.edu/yr312kt5378

Contains the device and metabolic data for the third *Static Training* participant.

#### Dataset S9: purl.stanford.edu/hq152jn6095

Contains the device and metabolic data for the fourth *Static Training* participant.

#### Dataset S10: purl.stanford.edu/mh986bj2257

Contains the device and metabolic data for the fifth *Static Training* participant.

#### Dataset S11: purl.stanford.edu/yb176ht8265

Contains the device and metabolic data for the first *Re-optimization* participant.

##### Dataset S12: purl.stanford.edu/sg494jd0004

Contains the device and metabolic data for the second *Re-optimization* participant.

##### Dataset S13: purl.stanford.edu/tr245wh8622

Contains the device and metabolic data for the third *Re-optimization* participant.

##### Dataset S14: purl.stanford.edu/jj186wr8258

Contains the device and metabolic data for the fourth *Re-optimization* participant.

##### Dataset S15: purl.stanford.edu/cb290rf2125

Contains the device and metabolic data for the fifth *Re-optimization* participant.
